# On the robustness to gene tree rooting (or lack thereof) of triplet-based species tree estimation methods

**DOI:** 10.1101/2024.11.22.624944

**Authors:** Tanjeem Azwad Zaman, Rabib Jahin Ibn Momin, Md. Shamsuzzoha Bayzid

## Abstract

Species tree estimation is frequently based on phylogenomic approaches that use multiple genes from throughout the genome. This process becomes particularly challenging due to gene tree heterogeneity (discordance), often resulting from Incomplete Lineage Sorting (ILS). Triplet and quartet-based approaches for species tree estimation have gained substantial attention as they are provably statistically consistent in the presence of ILS. However, unlike quartet-based methods, the limitation of rooted triplet-based methods in handling unrooted gene trees has restricted their adoption in the systematics community. Furthermore, since the induced triplet distribution in a gene tree depends on the placement of the root, the accuracy of triplet-based methods depends on the accuracy of gene tree rooting. Despite progress in developing methods for rooting unrooted gene trees, greatly understudied is the choice of rooting technique and downstream effects on species tree inference under realistic model conditions. This study involves rigorous empirical testing with different gene tree rooting approaches to establish a nuanced understanding of the impact of rooting on species tree accuracy. Moreover, we aim to investigate the conditions under which triplet-based methods provide more accurate species tree estimations than the widely-used quartet-based methods such as ASTRAL.

## 1 Introduction

The estimation of species trees using multiple loci has become increasingly common. However, combining multi-locus data is difficult, especially in the presence of gene tree discordance, where different genes may have different evolutionary histories. A traditional approach for species tree estimation from multiple genes is called concatenation (also called ‘combined analysis’), where alignments of the genes are concatenated into a supermatrix, which is then used to estimate the species tree. Although it is a widely used technique, concatenation can be problematic as it is agnostic to the topological differences among the gene trees, can be statistically inconsistent [46], and can return incorrect trees with high confidence [29, 16, 32, 11]. As a result, “summary methods”, which operate by computing gene trees from different loci and then combining the inferred gene trees into a species tree, are becoming increasingly popular [4].

Recent modeling and computational advances have produced summary methods that explicitly take gene tree discordance into account while estimating species trees from multi-locus data. While several biological processes, such as gene duplication and loss, incomplete lineage sorting (ILS), and horizontal gene transfer, can result in this gene tree conflict, ILS (modeled by the multi-species coalescent [27]), is potentially the most prevalent cause of gene tree heterogeneity. Therefore, recent literature has primarily focused on estimating species trees in the presence of ILS. Quartet and triplet-based summary methods [21, 28, 31, 43, 36, 33, 8, 52, 23] have gained significant attention as quartets (4-leaf unrooted trees) and triplets (3-leaf rooted trees) do not contain the “anomaly zone” [12–14], a condition where the most probable gene tree topology may differ from the species tree topology. ASTRAL [40, 41, 56], the most widely used summary method, is a quartet-based method that uses a dynamic programming (DP) approach to find a species tree that is consistent with the largest number of quartets induced by the set of gene trees. STELAR [23], on the other hand, is a triplet-based method which seeks a species tree by maximizing the number of consistent triplets with respect to the input gene trees. Both ASTRAL and STELAR are statistically consistent, have comparable accuracy and are scalable to large datasets. However, while ASTRAL has been a widely adopted choice, STELAR is not as popular as ASTRAL, possibly due to its limitation in analyzing unrooted gene trees. Gene trees are usually estimated using time reversible mutation models which make the root of the tree non-identifiable [14].

Unrooted gene trees are commonly transformed into rooted trees by incorporating an outgroup that places the root between the outgroup and the remaining taxa in the tree. Another approach involves introducing the assumption of a molecular clock. However, identifying a suitable outgroup proves to be challenging, and the use of a pre-specified outgroup may lead to biased root placement [42, 19]. Therefore, in the absence of a molecular clock or a reliable outgroup, alternative techniques for rooting phylogenetic trees have been developed [10, 44, 1, 6, 5, 50]. Despite the long history of thinking about tree rooting, there has been a lack of investigation into their impact in the context of estimating species trees using methods that rely on rooted gene trees.

In this study, we focus on triplet-based species tree estimating methods and investigate the robustness (or lack thereof) of these methods to variations in gene tree rooting. We report, on an extensive experimental study using a collection of simulated as well as empirical datasets, the performance of STELAR with gene trees rooted by different techniques. Furthermore, we identified different model conditions where STELAR with algorithmically rooted gene trees (i.e. rooted with methods other than outgroup rooting) outperformed not only STELAR paired with outgroup rooting, but also the state-of-the-art ASTRAL. These results indicate the potential for finding appropriate roots for the gene trees that result in better species tree estimations than ASTRAL, and thus we believe opens up the avenue for further research into the impact of gene tree rooting on the performance of triplet-based species tree estimation methods.

## 2 Experimental studies

### 2.1 Datasets

We studied a collection of previously used simulated and biological datasets to evaluate the impact of various rooting techniques on the triplet-based summary method STELAR. We also compared STELAR (with various rooting techniques) with ASTRAL, the leading coalescent-based summary method, which maximizes quartet-consistency and thus does not require rooted gene trees.

#### Simulated datasets

We used two biologically-based simulated datasets from [39], which were generated based on species trees estimated by MP-EST [33] on avian (48 taxa) and mammalian (37 taxa) datasets from [25] and [48], respectively. We used a 15-taxon dataset from [3] with a caterpillar-like (pectinate or ladder-like) model species tree. For 11 taxa [9], we specifically analyzed the regions exhibiting strong incomplete lineage sorting (ILS). We also analyzed two more relatively large datasets containing 200 and 500 taxa [41]. The simulated datasets we studied varied in many respects (number of genes, sequence length per gene, whether the sequence evolution is ultrametric or not, and the ILS level: low [2X], moderate [1X], and high [0.5X]). Thus, they represent a wide range of challenging model conditions on which we evaluated the impact of various rooting methods on the performance of the triplet-based species tree estimation method STELAR. Section 1 of the Supplementary Material provides more details in this regard.

#### Empirical datasets

##### Angiosperm dataset

We used the Angiosperm dataset of 310 nuclear genes analyzed in [54], [41]. The nuclear gene taxon sampling included 42 species representing all major angiosperm clades (35 families and 28 orders [20]). Three gymnosperms (*Picea glauca* [Moench] Voss, *Pinus taeda* L., and *Zamia vazquezii* D.W. Stev., Sabato & De Luca) and one lycophyte (*Selaginella moellendorffii* Hieron.) were included as outgroups. These three gymnosperms span the crown node of extant gymnosperms [55].

##### Amniota dataset

We used the Amniota dataset from [7] containing 16 species and 248 genes. We analyzed the nucleotide (nt) part of the dataset where *Xenopus tropicalis* and/or *Protopterus annectens* were used as outgroups.

### 2.2 Methods compared

We investigated the performance of different rooting techniques paired with STELAR, a statistically consistent triplet-based species tree estimation method.

There exist various rooting techniques for constructing and interpreting phylogenetic trees. The outgroup (OG) rooting method is the most commonly used technique for rooting phylogenetic trees. While this method generally provides better rooting accuracy when compared to other rooting methods [22], the main challenge lies in selecting an appropriate outgroup [53, 35, 47, 49, 38, 34]. In cases where an appropriate outgroup is unknown, inferring a root is possible using a molecular clock [22, 15]. Furthermore, there are rooting methods that take into account the distribution of branch lengths (and/or branch supports). Midpoint Rooting [17], Minimum Variance Rooting [37], and Minimal Ancestor Deviation [51] fall under this category. Other methods that do not root using an outgroup include gene duplication-based rooting [10, 24, 18], indel-based rooting [30, 44, 2], rooting species trees using distributions of unrooted gene trees [1], probabilistic co-estimation of gene and species trees [6], rooting using a non-reversible Markov model with the help of Multiple Sequence Alignments [5], and rooting species trees using the distribution of quintets induced by gene trees [50].

Thus, our experimental pipeline consists of the following steps: 1) take a set of unrooted gene trees, 2) root them using an appropriate rooting method of our choice, and finally, 3) infer the rooted species tree using a species tree estimation method (STELAR [23]). We used the following rooting methods: Outgroup Rooting (OG) [26], Midpoint Rooting (MP) [26], Minimum Variance rooting (MV) [37], Minimum Ancestor Deviation rooting (MAD) [51], both the “Search” (RD) and “Exhaustive” (RD-EX) modes of RootDigger [5], and Random rooting (RAND). We refer to STELAR paired with a particular rooting technique using the convention STELAR–⟨quartet-generation-technique⟩ (e.g, STELAR-OG, STELAR-MP, etc.). We also compared STELAR paired with different rooting techniques with ASTRAL (the latest ASTRAL-III version [56]), the leading species tree estimation method which maximizes quartet consistency. Further details of these rooting methods can be found in Section 2 of the Supplementary Material.

### 2.3 Evaluation Metrics

We compared the estimated trees (on simulated datasets) with the model species tree using normalized Robinson-Foulds (RF) distance [45], which is a widely used metric to measure tree error. The RF distance between two trees is the sum of the bipartitions (splits) induced by one tree but not by the other, and vice versa.

We also compared the triplet and quartet scores of candidate species trees. Triplet score of a rooted species tree *T* with respect to a particular set of rooted gene trees is defined as the number of triplets induced by the gene trees that the candidate tree *T* is consistent with. Maximizing triplet score is a statistically consistent criterion for estimating species trees [23]. We use the term true triplet score (TTS) when we compute the triplet score with respect to true gene trees (no estimation error). We similarly define and use the metrics quartet score and true quartet score, which consider quartet consistency instead.

To assess the correctness of the root placements inferred by various techniques, we computed the topological distance between the inferred root and the true root [5]. This is defined as the number of nodes in the path from the inferred/estimated root to a predefined true root. This distance quantifies the error in root placement without any scaling or normalization [5] based on tree size. This metric was introduced to quantify errors in root placement, addressing the limitations of the binary “percentage of correct rooting” measure, which failed to capture such nuances.

## 3 Results and discussion

### 3.1 Simulated Datasets

#### Results on 37-Taxa

Figure 1 shows the performance of STELAR with gene trees rooted by different methods and ASTRAL on the 37-taxon dataset under various model conditions. One of the key observations from the 37-taxon dataset is the robustness of the triplet-based species tree estimation method, STELAR, to gene trees rooted by various techniques. Most rooting methods, including MP, MV, and MAD, resulted in competitive species trees with no statistically significant differences (as seen from Figure 1). Surprisingly, while random rooting (RAND) performed worse in many instances, it still remained competitive under certain model conditions, such as 2X [low] ILS (Figure 1c), 1000 bp, and true gene trees (Figure 1b). Despite yielding comparable accuracy in the 2x ILS model condition (Figure 1c), STELAR performed substantially worse in all other model conditions when working with gene trees rooted by RD. On the other hand, RDEX rooting performed notably better in most model conditions.

**Fig. 1:**
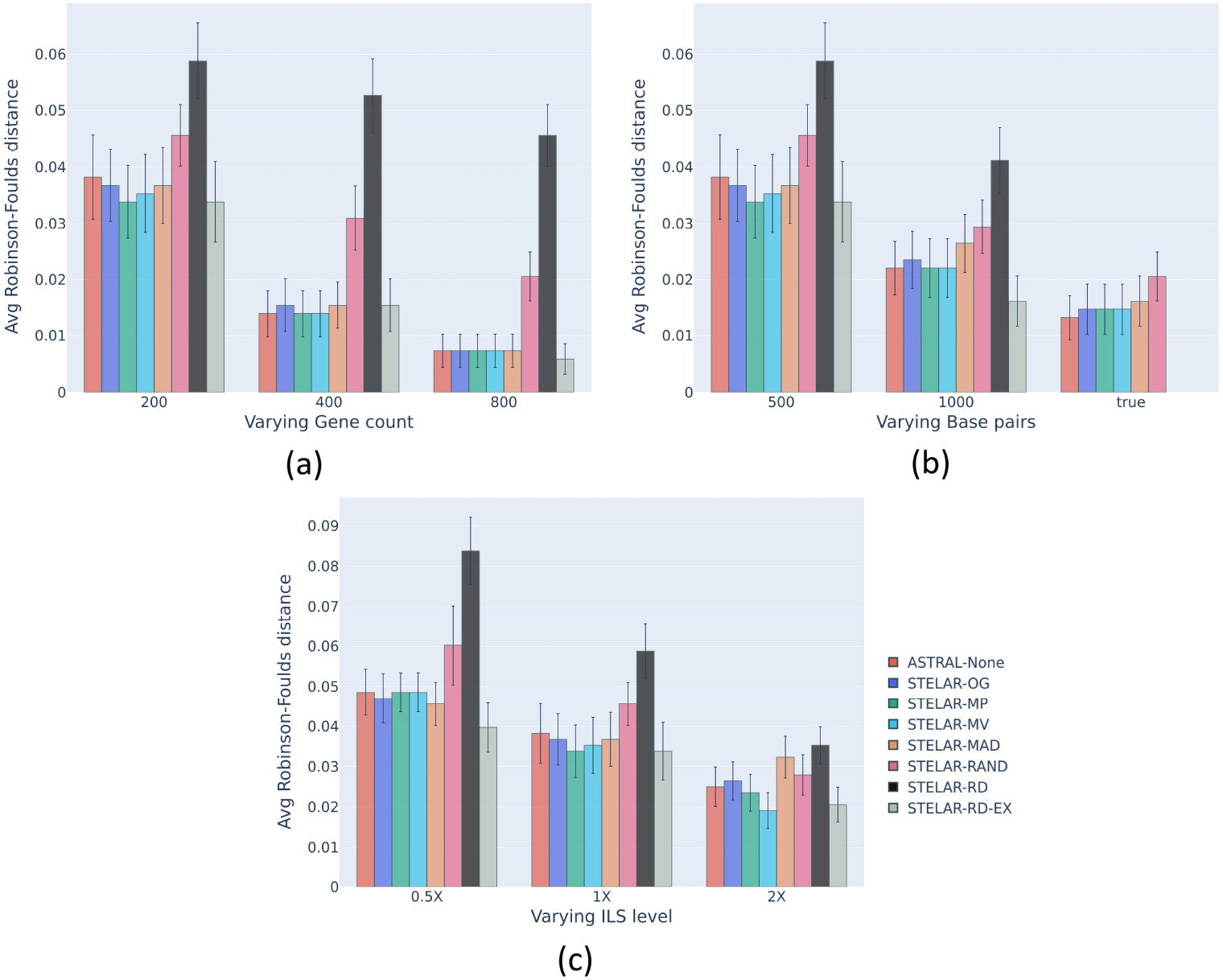
Comparison of Average Robinson-Foulds distance with standard error bars over 20 replicates for ASTRAL and STELAR with different rooting methods for the 37-taxa mammalian dataset. (a) We fixed the ILS at moderate (1X) and the base pairs at 500bp, and varied gene counts between 200, 400 and 800. RD could not be run on 400 genes, since the gene sequences corresponding to the gene trees were not available. We varied the sequence length, and consequently the gene tree estimation error, between 500, 1000 and true-gt while keeping moderate ILS (1X) and 200 gene trees. (c) We set the gene trees at 200, the sequence length at 500bp, and varied the ILS between high (0.5X), moderate (1X) and low (2X)

Interestingly, STELAR sometimes yielded more accurate species trees with rootings other than the known/true OG. There were model conditions where gene trees rooted using MP, MV, MAD, and RD-EX resulted in superior species trees compared to OG and even ASTRAL, albeit these differences were not statistically significant. Such cases include MAD in 0.5X [high] and 1X [moderate] ILS cases, MP, MV in the 1X-200gene-500bp and 2X-200gene-500bp model conditions, and RD-EX in all conditions except 1x-400gene-500bp.

The general patterns were consistent with our expectations: for all methods, the species tree estimation accuracy was improved by increasing the number of genes and sequence lengths (i.e., decreasing the gene tree estimation error), but was reduced by increasing the amount of gene tree discordance (i.e., the amount of ILS). In general, the choice of rooting techniques becomes less impactful in “easier” model conditions (e.g., higher gene count and more base pairs), with STELAR with MP, MV, OG, and MAD all yielding the same or similar tree accuracy (e.g., under conditions with 800 genes and true gene trees).

Next, we compared the relative performance of different rooting techniques in terms of species tree accuracy. RD-EX, MP, and MV perform notably well on this dataset, especially in “unfavorable” model conditions with fewer genes, shorter gene sequences, and higher ILS. MP and RD-EX matched or outperformed most other methods, including ASTRAL, across various model conditions. RD-EX especially performed well in conditions with high (0.5X) ILS and high basepair count (1000bp). Similarly, MV performed on par with other methods, including ASTRAL, and showed significant improvements under low ILS (2X) conditions. Although MAD achieved one of the best species tree accuracies at high ILS (0.5x), its performance remained largely unaffected by varying levels of ILS. This unresponsiveness of MAD was also observed toward varying gene tree estimation errors (controlled by sequence lengths). However, like other methods, MAD’s performance improved with an increasing number of genes.

To further investigate why RD-EX, MP, MV, and occasionally MAD yielded better RF scores than OG, and why RD performed poorly, we conducted a series of experiments examining triplet and quartet scores, as well as the correctness of the roots of the gene trees and species trees inferred by various techniques. First, we compared the triplet scores (Table 1 and Supplementary Table S2 for higher precision) and quartet scores (Supplementary Table S4) of the estimated species trees. As expected, due to the statistical consistency property of the triplet score, in almost all cases involving STELAR, a lower RF score corresponds to a higher triplet score. For instance, under high ILS conditions (0.5X), MAD achieved the highest triplet score and the lowest RF score. Interestingly, OG, which is expected to yield the highest triplet score when the outgroup is known for simulated datasets, did not perform as expected under certain model conditions (e.g., 1X-200gt-1000bp and 1X-400gt-500bp). In these scenarios, STELAR using gene trees rooted by OG resulted in marginally lower triplet scores and higher RF scores compared to STELAR with gene trees rooted by MP and MV. There were a few exceptions to this trend, such as in the 1X-200gt-500bp and 2X-200gt-500bp conditions, where OG achieved higher triplet scores despite MP and MV performing better in terms of RF scores. In some cases (e.g 0.5X-200gt-500bp and 1X-200gt-1000bp) where RD-EX achieved the best RF scores, it displayed slightly lower triplet scores - an exceptional trend that can be attributed to higher variance in the RF scores that were averaged.

**Table 1:** Average Triplet Score (TS) (in hundreds)

| Dataset | Model Cond<sup>n</sup> | ASTRAL | STELAR |  |  |  |  |  |  |
| --- | --- | --- | --- | --- | --- | --- | --- | --- | --- |
|  |  |  | MAD | MP | MV | OG | RAND | RD | RD-EX |
| 15-taxon | 100-100 | 152 | 172 | **177** | **177** | **177** | 165 | 177 | 169 |
|  | 100-1000 | 173 | 201 | 203 | 203 | 203 | 185 | **203** | 198 |
|  | 100-true | 259 |  | 240 | 222 | **328** | 199 |  |  |
|  | 1000-100 | 1499 | 1716 | **1759** | **1759** | **1759** | 1638 | 1759 | 1679 |
|  | 1000-1000 | 1702 | 1987 | 1996 | 1996 | 1996 | 1823 | **2018** | 1948 |
|  | 1000-true | 2588 |  | 2411 | 2186 | **3276** | 2038 |  |  |
| 37-taxon | 0.5X-200-500 | 9285 | **9895** | 9895 | 9891 | 9893 | 8741 | 8431 | 9818 |
|  | 1X-200-1000 | 10511 | 11584 | **11585** | **11585** | 11585 | 9613 | 9401 | 11382 |
|  | 1X-200-500 | 9814 | 11014 | 11017 | 11016 | **11024** | 9527 | 9325 | 10940 |
|  | 1X-200-true | 13059 | **14107** | **14107** | **14107** | **14107** | 9988 |  |  |
|  | 1X-400-500 | 20013 | 22027 | **22027** | **22027** | 22027 | 19056 | 19051 | 21846 |
|  | 1X-800-500 | 40046 | **44160** | **44160** | **44160** | **44160** | 38107 | 38097 | 43763 |
|  | 2X-200-500 | 10062 | 11529 | 11548 | 11540 | **11549** | 9890 | 9670 | 11456 |
| 200-taxon | estimated | 7171596 | 7553494 | 7652796 | 7497983 | 7596758 | **7858044** |  |  |
| Amniota | Amino Acid | 251 |  | 308 | 251 | **381** | 298 |  |  |
|  | Nucleotide | 312 |  | 346 | 284 | **424** | 284 |  |  |
| Angiosperm | Nuclear | 9127 | **12146** | 12007 | 12007 | 12007 | 8679 |  |  |

We also looked at the triplet score of ASTRAL-estimated trees. Though the ASTRAL output species tree is meant to be unrooted, we use the root implied from its Newick representation. Expectedly, ASTRAL–which maximizes quartet scores–consistently achieved substantially lower triplet scores than STELAR despite its higher accuracy. ASTRAL often failed to achieve the best RF scores, even though it always had the highest quartet scores (see 0.5x-200gt-500bp, 1x-200gt-500bp, 2x-200gt-500bp model conditions). Similar to triplet-scores, when STELAR-MP and STELAR-MV outperformed STELAR-OG in species tree accuracy (e.g., 1X-200gt-1000bp and 1X-400gt-500bp), they tended to achieve higher quartet scores. However, as with triplet scores, there were exceptions to this trend in some model conditions such as 0.5x-200gt-500bp and 2X-200gt-500bp, and for the few aforementioned cases of RD-EX.

Next, we examined the percentage of correctly rooted gene trees among the inputs to STELAR for each rooting method. This simulated dataset is rooted using the outgroup (Chicken, Turkey). Aside from the obvious 100% accuracy from OG, we see MV consistently reaching the second best, closely followed by MP and then MAD (Supplementary Table S6). RAND, RD, and RD-EX scored poorly in this metric. To further evaluate gene tree rooting accuracy for the rooting methods, we calculated the average topological distance between the true root and inferred root for the gene trees, in Table 2. This analysis supports the trend seen above: by definition, OG scores 0, indicating perfect accuracy, while MV, MP, and MAD demonstrate progressively increasing topological distances, reflecting their respective accuracies in inferring the root position. RDEX, RAND, and RD give higher distances on average, corroborating their low percentage of correct rootings.

**Table 2:** Avg. Topological Distance of estimated root from true root of gene trees.

| Dataset | Model Cond<sup>n</sup> | MAD | MP | MV | OG | RAND | RD | RD-EX |
| --- | --- | --- | --- | --- | --- | --- | --- | --- |
| **15-taxon** | 100gt-100bp | 3.54 | 0.00 | 0.00 | 0.00 | 5.25 | 3.69 | 5.34 |
|  | 100gt-1000bp | 0.76 | 0.00 | 0.00 | 0.00 | 5.03 | 3.37 | 4.77 |
|  | 100gt-true |  | 2.95 | 3.16 | 0.00 | 5.05 |  |  |
|  | 1000gt-100bp | 3.52 | 0.00 | 0.01 | 0.00 | 5.33 | 3.74 | 5.27 |
|  | 1000gt-1000bp | 0.82 | 0.00 | 0.00 | 0.00 | 5.13 | 3.44 | 4.81 |
|  | 1000gt-true |  | 2.92 | 3.11 | 0.00 | 4.96 |  |  |
| **37-taxon** | 0.5X-200-500 | 1.42 | 0.63 | 0.55 | 0.00 | 7.61 | 8.09 | 4.38 |
|  | 1X-200-1000 | 0.60 | 0.39 | 0.28 | 0.00 | 7.65 | 8.07 | 4.19 |
|  | 1X-200-500 | 1.37 | 0.56 | 0.44 | 0.00 | 7.75 | 8.15 | 4.22 |
|  | 1X-400-500 | 1.34 | 0.55 | 0.44 | 0.00 | 7.65 | 8.17 | 4.22 |
|  | 1X-800-500 | 1.35 | 0.56 | 0.45 | 0.00 | 7.70 | 8.17 | 4.25 |
|  | 2X-200-500 | 1.30 | 0.56 | 0.44 | 0.00 | 7.69 | 8.11 | 4.18 |
| **200-taxon** | estimated | 4.68 | 4.74 | 4.76 | 0.00 | 11.09 |  |  |
| **500-taxon** | true | 2.63 | 2.51 | 2.33 | 0.00 | 11.74 |  |  |
|  | estimated | 5.61 | 5.92 | 5.88 | 0.00 | 14.15 |  |  |

Despite the fact that MP and MV did not achieve the same level of rooting accuracy as OG, as previously noted, the gene trees rooted by MP and MV produced species trees that were equally good or even better than those inferred using OG. This indicates the robustness of triplet-based methods in species tree estimation from gene trees rooted using different techniques under certain model conditions. RAND, as expected, failed to recover the correct root in most cases (around 98% of the time). Surprisingly, RD was even worse than random rooting (RAND), almost never recovering the correct root. This poor rooting accuracy of the gene trees rooted by RD likely explains the poor performance of STELARRD. STELAR-RD-EX, despite performing well in terms of RF scores in most cases, yielded low gene-tree rooting accuracy. Such a trend is better analyzed in the next dataset.

Finally, we examined the root of the estimated species trees. Unlike the quartet-based species tree estimation method ASTRAL, triplet-based methods work with rooted gene trees and infer a rooted species tree that maximizes the triplet score, making the rooting implied by the output of STELAR meaningful. We, therefore, analyzed the percentage of correct rooting in the inferred species trees as well as the average topological distance between the inferred species tree roots and the true root, as shown in Supplementary Tables S7 and S9 respectively. Interestingly, despite the previously noted differences in RF scores, triplet scores, and quartet scores, all triplet-based methods except for STELARRD, STELAR-RD-EX, and STELAR-RAND correctly predicted the rooting of the species tree across all model conditions. Moreover, the topological distances between the roots inferred by RD-EX, RAND, and RD variants of STELAR and the true roots were not substantially large. These findings highlight the robustness of the triplet-based species tree estimation method STELAR in accurately inferring the root of the estimated species trees, even when the roots of the input gene trees differ.

#### Results on 15-Taxa

A general trend similar to that in the 37-taxa dataset is also observed in the analysis of the 15-taxa dataset: all methods, including STELAR with different gene tree rootings, yield better performances with “easier” model conditions. However, a primary point of interest in this dataset is the significantly better performance of STELAR-MAD in terms of RF scores (Figure 2), even outperforming ASTRAL in all model conditions ^1^ -a more prominent variation of RD-EX’s performance in the 37 taxa dataset. Despite displaying lower Triplet Scores compared to the other STELAR variants (Table 1), and lower Quartet scores compared to ASTRAL (Supplementary Table S4), STELAR-MAD showed better robustness to gene tree estimation errors as reflected in its mostly unmatched True Triplet Scores (Table 3 and for higher precision, Supplementary Table S3) and True Quartet Scores (Supplementary Table S5). Referring to the average topological distance (Table 2) we note that, compared to the other rooting methods, MAD rooted the gene trees further from the true root. This seemingly “inaccurate” rooting by MAD actually helped STELAR in overcoming gene tree estimation errors, resulting in the best performances in this dataset.

**Table 3:** True Triplet Scores (in hundreds)

| Dataset | Model Cond<sup>n</sup> | ASTRAL | STELAR |  |  |  |  |  |  |
| --- | --- | --- | --- | --- | --- | --- | --- | --- | --- |
|  |  |  | MAD | MP | MV | OG | RAND | RD | RD-EX |
| 15-taxon | 100gt-100bp | 196 | 300 | **305** | **305** | **305** | 202 | 282 | 215 |
|  | 100gt-1000bp | 217 | **328** | 325 | 325 | 325 | 209 | 263 | 238 |
|  | 1000gt-100bp | 1875 | **3210** | 3092 | 3092 | 3092 | 1981 | 2897 | 2179 |
|  | 1000gt-1000bp | 2316 | **3276** | 3268 | 3268 | 3268 | 1992 | 2628 | 2319 |
| 37-taxon | 0.5X-200-500 | 11057 | 12481 | 12481 | **12482** | 12481 | 9208 | 9204 | 11497 |
|  | 1X-200-1000 | 12013 | 14104 | **14107** | **14107** | 14107 | 9987 | 8330 | 12815 |
|  | 1X-200-500 | 11452 | 14104 | **14105** | 14104 | 14095 | 9981 | 8697 | 12918 |
|  | 1X-400-500 | 23559 | 28201 | **28202** | **28202** | 28202 | 19966 | 19959 | 25627 |
|  | 1X-800-500 | 46006 | **54795** | **54795** | **54795** | **54795** | 38810 | 33712 | 49786 |
|  | 2X-200-500 | 11743 | 14974 | **14975** | 14990 | 14965 | 10348 | 10343 | 13574 |

**Fig. 2:**
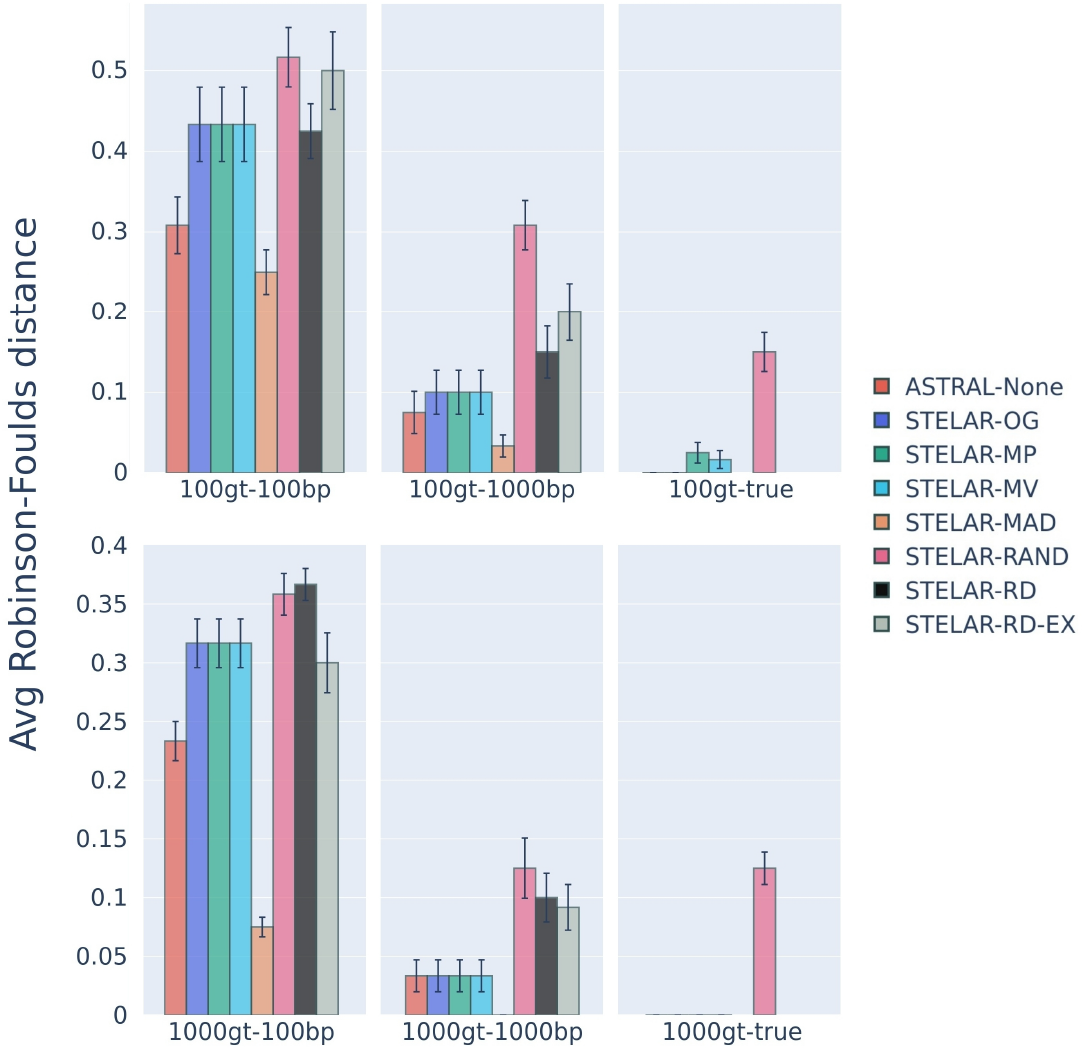
Comparison of Robinson-Foulds distance for ASTRAL and STELAR with different rooting methods on the 15-taxa dataset. We varied the number of estimated gene trees (100genes - 1000genes) as well as the sequence length (100bp - 1000bp). We also analyzed model conditions with true gene trees; however, due to absence of branch lengths, we could not run MAD on these cases (100-true and 1000-true).

Aside from the exceptional performance by STELAR-MAD, we find that the MP and MV variants of STELAR infer the same species tree as the OG variant in most cases, attesting to the robustness of STELAR to gene tree rooting. These variants also yield competitive species tree accuracies with respect to ASTRAL, aside from the “difficult” model conditions of lower base-pair counts (100bp). Notably, STELAR-RAND yields signifcantly worse performances in “easier” conditions–a trend not observed in the 37 taxa dataset. This indicates a possible correlation between STELAR-RAND’s performance and the number of taxa under consideration. Aside from the exception of STELAR-MAD, quartet scores (Supplementary Table S4) were better correlated to species tree accuracy (Figure 2) compared to triplet scores (Figure 1) in this dataset. Furthermore, similar to the 37 taxa dataset, we notice high proficiency from STELAR in correctly predicting the species tree root (Supplementary Table S7) despite inaccuracies in the input gene tree rootings (Supplementary Table S6).

#### Results on relatively large number of taxa

On the 200 taxa dataset, STELAR-OG outperformed all other variants of STELAR in terms of RF scores by a significant margin, as seen from Figure 3(a). Strikingly, RAND showed a performance comparable to that of MP, MV, MAD. This also translated to the triplet scores, where RAND yielded the highest triplet score among all methods. Though ASTRAL yielded slightly better performance when compared to STELAR-OG, this was not statistically significant.

**Fig. 3:**
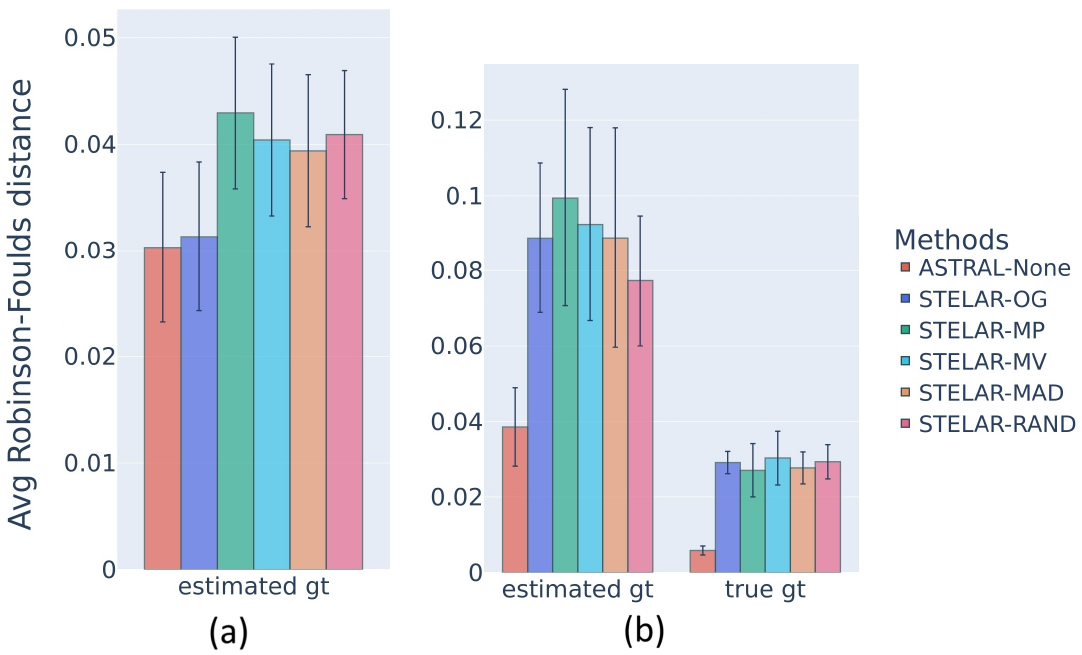
RF Scores for higher-order taxa datasets. (a) 200 Taxa (b) 500 taxa

A similar robustness to gene tree rooting for STELAR is noticed in the 500 taxa dataset depicted in Figure 3(b). In the simulated case, STELARRAND came in second, outperforming even STELAR-OG in terms of RF scores. STELAR-RAND also showed a performance comparable to the other STELAR variants in the true gene tree case. However, ASTRAL displayed statistically significant improvement in terms of RF scores when compared to the STELAR variants.

The number of species trees in the search space that differs from the “best” tree by *k* edges increases as the number of taxa increases. As such, STELAR has more options to arrive at a species tree with a better/same RF score, despite working on randomly rooted gene trees. Furthermore, a single mismatched bipartition has a more severe impact on the RF score when the number of taxa is lower. The better performance of STELAR-RAND in terms of RF score for large datasets can quite possibly be attributed to these facts. Thus, STELAR grows increasingly robust to gene tree rooting as the number of taxa increases.

### 3.2 Biological Datasets

#### Results on Angiosperm Dataset

Analyses on this dataset aim to address long-standing questions in the phylogeny of Angiosperms, with the key challenges lying in the phylogenetic placement of *Amborella trichopoda* and its relative branching order with other lineages like Nymphaeales (i.e. *Nuphar* or waterlilies)

STELAR-MAD correctly placed *Amborella* as a sister to the outgroup comprised of (Selaginella, (Zamia, (Pinus and Picea))) (Fig. 4b). As such, the tree inferred by this method correctly places Amborella as the sister to the other Angiosperms. But, this method could not predict Nuphar as the next sister to the angiosperms (as inferred in [54], nor does it predict the clade of (Amborella, Nuphar) as done in [41]. These relationships were not inferred by the other STELAR methods as well.

**Fig. 4:**
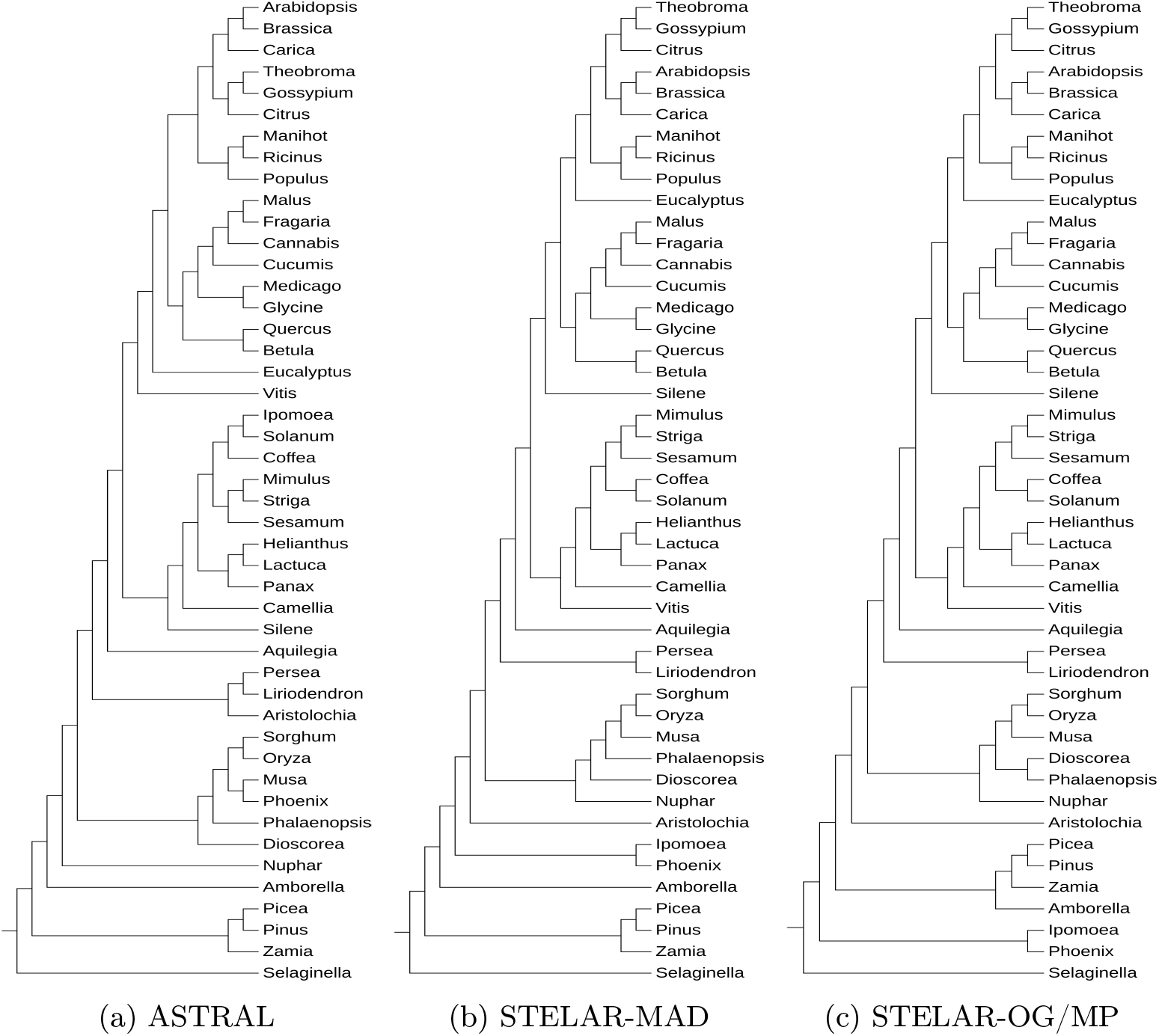
Estimated Species trees for Angiosperm dataset

Furthermore, unlike STELAR-MAD, the OG, MP, and MV variants of STELAR misplaced *Selaginella*, causing *Amborella* to be placed as a sister to the clade comprised of the 3 other outgroup taxa (Zamia, (Pinus, Picea)) (Fig. 4c). All STELAR variants had issues with the following placements: swapped positions of Vitis and Silene, misplaced Eucalyptus, Phoenix, Ipomoea and Aristolochia. The species tree from ASTRAL is the same as stated in [55], and does not misplace the aforementioned taxa (Fig. 4a). It predicts the clade of (Amborella, Nuphar) as sister to the other Angiosperms.

It is worth noting that none of the methods originally rooted the estimated species trees at the outgroup. The STELAR methods predicted the root at (Phoenix, Ipomoea) and ASTRAL originally rooted at *Camellia*. For ease of discussion, the species trees were all rerooted at *Selaginella*.

#### Results on Amniota Dataset

We analyzed the nuleotide (DNA) variant of the dataset from [7], containing 16 amniota taxa. The key challenge lies in resolving the position of turtles relative to crocodiles and birds. Past studies like [36] place turtles as sisters to the archosaurs clade (comprised of birds and crocodiles).

Both the MP and MV variants of STELAR correctly predicted the Archosaurs clade, similar to ASTRAL (fig. 5a). However, STELAR-OG failed to predict the Archosaurs clade as seen from fig 5b. All methods correctly predicted the other significant clades like turtles, crocodiles, birds, and squamates.

**Fig. 5:**
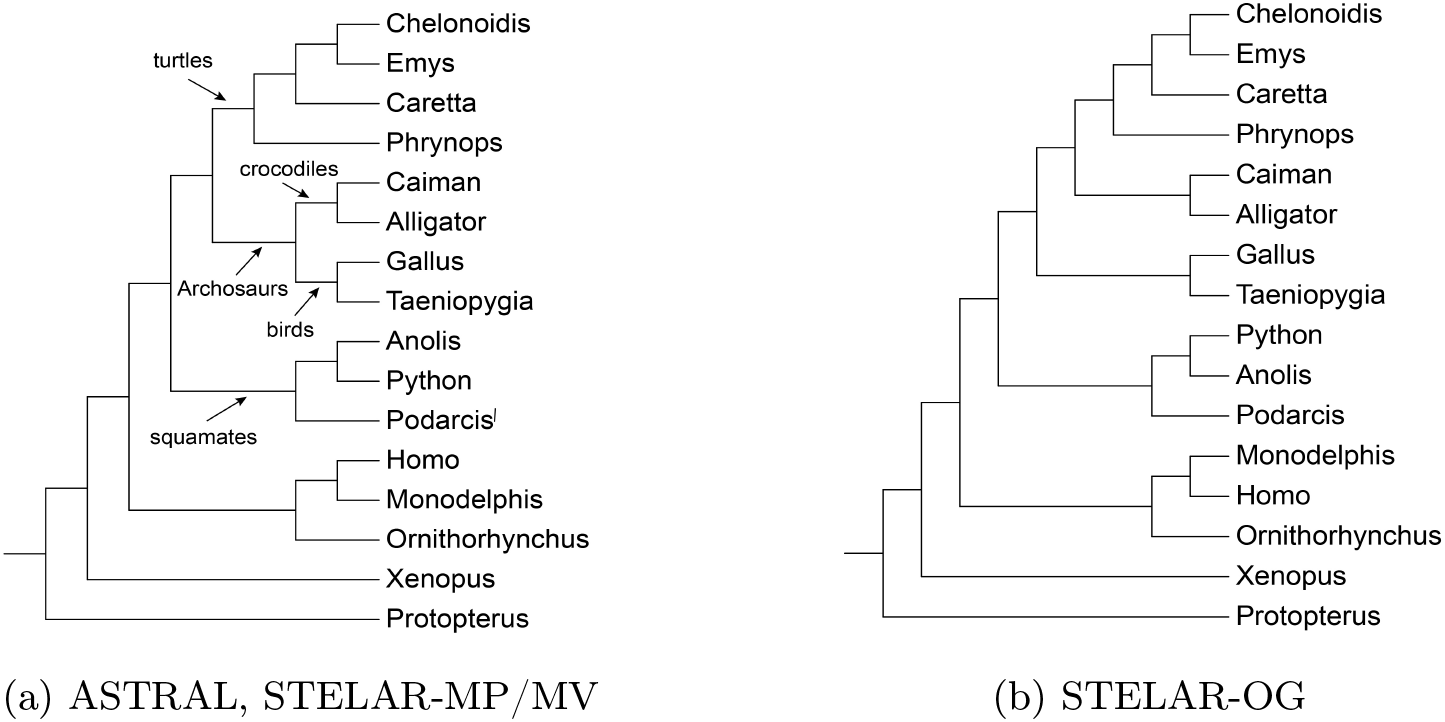
Estimated species trees for Amniota (Nucleotide) dataset

It is worth noting that only STELAR-OG could correctly predict the rooting at *Protopterus*; ASTRAL and the other STELAR variants could not, and thus yielded lower triplet scores (Table 1) despite their correct clade predictions. The rooting inferred from ASTRAL is *Ornithorhynchus*, whereas STELAR-MV placed the root at the clade ((Protopterus, Xenopus), (Ornithorhyncus, (Homo, Monodelphis)))

## 4 Conclusion

Both quartets and triplets avoid the “anomaly zone” – a scenario where the most likely gene tree topology may differ from the species tree topology [12–14]. Thus, statistically consistent methods have been developed by maximizing quartet and triplet consistency. While quartet-based summary methods like ASTRAL are in wide use, triplet-based methods like STELAR has not gained notable attention from the community. Moreover, triplet-based methods require rooted gene trees which is often difficult to obtain in the absence of molecular clock or reliable outgroups. Consequently, various techniques to root a given set of unrooted gene trees have been developed, differing in the type of data that can be analyzed, the assumptions about the evolutionary dynamics of the data, and their scalability or general applicability. However, little is known about the impact of these rooting techniques on species tree inference.

In this study, we considered a broad range of rooting approaches and their effects on species tree inference by maximizing triplet consistency. We found that STELAR was robust to the choice of rooting in most simulated model conditions as well as on real biological datasets (e.g., angiosperm dataset); where gene trees with algorithmic rootings performed competitively when compared to those rooted at the outgroup. The effect of gene tree rooting on species tree accuracy diminished with increasing taxa, to an extent where random rooting performed on par with the other methods in 200 and 500 taxa datasets. Randomness in rooting also seemed to have less impact in “more difficult” model conditions (higher ILS, lower basepairs, lower gene counts). These observations attest to the acceptability of STELAR in datasets where the outgroup is difficult to infer.

In addition, our studies also revealed conditions that yielded notable differences in species tree accuracy based on different gene tree rooting techniques. More interestingly, under some model conditions (like 0.5X-200-500, 1X-200-500, 2X-200-500, and 1X-200-1000 for 37 taxa and almost all cases for 15 taxa), STELAR produced more accurate species trees with rootings other than at the known outgroup. In the cases mentioned above, STELAR with algorithmic rootings also outperformed the state-of-the-art species tree estimation methods ASTRAL in terms of accuracy, suggesting that ‘correct’ rootings can lead to significant improvements in performance for STELAR.

Furthermore, we investigated triplet and quartet scores to better understand the effect of gene tree rooting on the performance of STELAR. Both of these metrics are statistically significant, but under practical model conditions with limited numbers of genes with gene tree estimation errors, achieving the highest quartet scores (as done by ASTRAL) or the highest triplet score (STELAR-OG) does not necessarily correspond to the best RF score, as seen in many model conditions. However, throughout our experimentation, we observe that quartet scores tend to be more correlated with the RF score than triplet scores.

To estimate the error in rooting, we calculate - for each method - the percentage of “correct” roots (w.r.t the true root) both for the input gene trees and for the species trees inferred by STELAR. Our study shows that STELAR is quite adept at predicting the correct root for the output species tree, even when the input gene trees have substantially different rootings (as clearly seen in the 37 taxa dataset). To better quantify rooting error, we introduce the metric of topological distance of the inferred root from the true root, which provides valuable insights into the effect of rooting on STELAR’s performance.

One such significant insight is the ability of STELAR to be robust to gene tree estimation error when provided appropriately rooted gene trees as input. We affirm this by calculating the triplet and quartet scores of STELAR outputs with respect to the true gene trees (gene trees free from all forms of estimation errors) - naming these metrics as true triplet scores and true quartet scores respectively. With increasing gene tree estimation error, maximizing the triplet score by rooting at/near the outgroup will not necessarily yield species trees with the best RF score. Certain algorithmic rooting methods can figure out roots that reduce the effect of this gene tree estimation error on STELAR’s performance. A noteworthy instance of this phenomena is the performance of STELAR-MAD in the 15-taxa dataset, where a seemingly “uncommon” root further from the true root helped it perform significantly better than the other methods.

To test whether the robustness of STELAR carries over to real-world biological datasets, we ran experiments on two empirical datasets. We found that STELAR shows similar robustness to gene tree rooting, as it yielded similar performances with algorithmically rooted inputs when compared to gene trees rooted at the outgroup. Furthermore, STELAR when paired with algorithmic rootings succeeded in predicting some important clades (like the Archosaurs clade in the Amniota-NT dataset), which was not the case for STELAR-OG.

Despite our thorough experimentation across diverse datasets and model conditions, we acknowledge limitations and potential areas for extension. Firstly, we explored datasets where the primary source of gene tree discordance was from Incomplete Lineage Sorting (ILS); this study can be extended to datasets where discordance arises from Horizontal Gene Transfer as well as gene duplication and loss. Furthermore, given the mixed performance of various rooting techniques, relative performance on finite data clearly depends on the model conditions. Hence, we do not make any general recommendation in favor of one type or rooting over another. And although we observe certain strengths for each method (eg. low ILS conditions for MV, high ILS for RD-EX, generally good performance for MP, high gene tree estimation error for MAD), extending our experiment to diverse datasets with finer control over the dataset generation parameters (like true species tree topology, varying speciation rates, controlled rates of gene tree estimation errors) will help us further pinpoint the strengths and weaknesses of each method. Moreover, the preprocessing steps for the rooting methods can be improved: for example, MP, MV and MAD rooting methods all require, and are heavily influenced by, branch lengths. Thus in datasets without branch lengths, we set each branch to have unit length, which sometimes adversely affected STELAR’s performance– eg MP, MV variants in 48 taxa (details in Section 3.3 of the Supplementary Material) and the 100gene-true model condition of the 15 taxa dataset. This aspect can be improved by using better branch length estimation algorithms. Furthermore, it was not possible to note the quartet and triplet Scores (using the functionalities provided by the ASTRAL and STELAR softwares respectively) for large datasets (e.g. 500 taxa) due to overflow errors.

## Footnotes

1 except for 100gt-true and 1000gt-true, since the branch lengths were absent in these cases and the MAD rooting method cannot be run on such gene trees

