## Supplementary Materials for "On the robustness to gene tree rooting (or lack thereof) of triplet-based species tree estimation methods"

#### 1 Dataset

##### 1.1 Simulated Dataset

We used two biologically-based simulated datasets from [22], which were generated based on species trees estimated by MP-EST [16] on avian and mammalian datasets from [14] and [28], respectively. For the avian simulations, gene sequence lengths were varied (250bp, 500bp, 1000bp, and 1500bp), while in the mammalian simulations, we examined sequence lengths of 250bp, 500bp, and 1000bp to assess the impact of phylogenetic signal. In both simulations, we modeled three levels of incomplete lineage sorting (ILS) by scaling internal branch lengths in the species tree and also explored varying gene counts.

15-taxon dataset was taken from [2] with a caterpillar-like (pectinate or ladder-like) model species tree, featuring 12 consecutive short internal branches (0.1 coalescent units) – creating conditions for high levels of ILS. Ultrametric gene trees were simulated along this tree under a multi-species coalescent model, adhering to a strict molecular clock without branch length transformations. Sequence data were generated for each gene tree, and four model conditions were constructed by using gene sequence lengths of 100 or 1000 sites and using either 100 or 1000 genes.

For 11 taxa, we specifically analyzed the regions exhibiting strong incomplete lineage sorting (ILS). The number of genes analyzed varied incrementally, with counts set at 5, 15, 25, 50, and 100 genes.

The simulated datasets we studied varied in many respects (number of genes, sequence length per gene, whether the sequence evolution is ultrametric or not, and the ILS level). Thus, they represent a wide range of model conditions on which we evaluated the impact of various rooting techniques on the performance of triplet-based species tree estimation method STELAR.

Table S1: Properties of the simulated datasets. The level of ILS is represented by the average topological distance between the true gene trees and the true species tree.

| Dataset | ILS level | # genes | # sites | # Replicates | Ref. |
|---|---|---|---|---|---|
| 11-taxon | 85% | 5 - 100 | 2000 | 20 | [6] |
| 15-taxon | 82% | 100 - 1000 | 100 - 1000 | 10 | [2] |
| 37-taxon | 18%, 32%, 54% | 200 - 800 | 500 -1000 | 20 | [22] |
| 48-taxon | 35%, 47%, 59% | 25 - 1000 | 250 -1000 | 20 | [22] |

We also analyzed two more relatively large datasets containing 200 and 500 taxa. For both 200-taxa and 500 taxa datasets, we analyzed tree lengths 2 M generations with speciation rates 1e-6 per generation. SimPhy was used to simulate species trees [23] based on the Yule process, defined by parameters such as the number of taxa, the maximum tree length, and the speciation rate, which together specify a model condition.

The tree length impacts the amount of ILS, with lower length resulting in shorter branches, and therefore higher levels of ILS. We used medium tree length (2 M) that indicates moderate level ILS. 10 replicates of each dataset (200 taxa, 500 taxa) were analyzed, with each replicate comprising 1,000 gene trees. These datasets were simulated according to the multi-species coalescent model with the population size fixed to 200000 [23].

#### 2 Rooting Methods Compared

Phylogenetic trees can be categorized as rooted or unrooted, each serving distinct purposes in evolutionary research. Rooted trees identify the last common ancestor, providing crucial insights into the directionality of evolution, whereas unrooted trees focus on the relationships among taxa without specifying evolutionary paths. Many methods, each with their own strengths and drawbacks, have been established for rooting unrooted phylogenetic trees. The rooting methods are described below:

##### 2.1 Outgroup Rooting Method (OG)

The outgroup method is a popular rooting approach in phylogenetics [4,11,13,32,36], often used to determine the evolutionary starting point of a tree. This method assumes that one or more taxa (the outgroup) are divergent from the main group being studied (the ingroup). The branch connecting the outgroup and ingroup sets the evolutionary baseline for the tree [5, 37]. The outgroup method not only aids in understanding ingroup evolution but also helps in identifying unique features within ingroup sequences. However, selecting an appropriate outgroup is challenging [17, 18, 21, 27, 30, 34], particularly for datasets with a large number of taxa where a consensus about the outgroup is lacking [24]. If an outgroup is too distantly related to the ingroups, it might adversely impact the rooting accuracy owing to a substantially different molecular evolution. Conversely, outgroups that are too closely related with the ingroups may fail to perform its functions as an appropriate outgroup.

Incorrect outgroup selection can also lead to long branch attraction (LBA), a phenomenon where distant outgroup taxa erroneously influence the tree due to their large divergence time or rapid evolution [32]. This results in artifactual or random rooting [10, 35] . To mitigate LBA, several criteria, such as low substitution rate and phylogenetic proximity, are suggested for choosing outgroups, particularly in complex cases like arthropod classes [25]. Additionally, [9] recommend using multiple outgroup samples within the sister group to minimize LBA and enhance the robustness of the outgroup rooting method. Gatesy et al. [8] demonstrated that introducing just one additional taxon can significantly alter the resulting tree topology, impacting even the existing ingroup taxa. Holland's simulations further confirm that outgroups that influence ingroup topologies are common [12].

OGs can also be used as "true" roots, if the original root is known beforehand (as in the case in simulated datasets).

##### 2.2 Mid-point Rooting Method (MP)

Midpoint rooting in phylogenetics is a method where the root is placed at the midpoint between the two furthest tips of the tree. [7,15] This method works effectively if the tree exhibits constant rates of evolution, relying on the assumption of a molecular clock and homogenous evolution rates across branches [12]. Midpoint rooting is particularly suitable for balanced trees but has limitations, especially if the tree data is not clocklike or the topology is unbalanced. The midpoint rooting method usually displays an impressively high success rate, which is especially remarkable in the multiple outgroup, consistently rooted datasets(MOC). [11]

This technique is frequently used in studies where outgroups are not available, such as in viral genetics . An example is [29], who applied midpoint rooting to analyze the evolutionary relationships of SARS coronaviruses, focusing on genes encoding envelope matrix and nucleocapsid proteins. In [20], phylogenetic trees were rooted using Midpoint-rooting method which helped in genome-based phylogeny and classification of prokaryotic viruses. Moreover, phylogenetic reconstuction was done in [38] using midpoint rooting technique. The appropriateness of midpoint rooting is supported by the results of Tajima's relative rate test [31], indicating no rate heterogeneity among the coronavirus groups [29]. It is advised to not use MP as the default method and to only use it if the outgroup method is not applicable e.g. due to a lack of priori knowledge regarding the outgroup or LBA [11].

##### 2.3 Minimum Variance Rooting Method (MV)

This section discusses the rooting method introduced in [19], which focuses on reducing the variance of root-to-tip distances in the phylogenetic tree. It is effective in cases where deviations from the strict molecular clock are random and can be implemented in a linear time algorithm, similar to the traditional midpoint rerooting method.

The effectiveness of this method has been explored through extensive simulations, considering factors like gene tree estimation error, divergence from the clock, and outgroup distance to ingroups. [19] showed that MV rooting performs better or at least as well as midpoint rooting across various conditions, especially when the divergence from the clock is less and outgroup distance is smaller. In [1], this method was used to root phylogenies under Outgroup-free rooting.

##### 2.4 Root-Digger Rooting Method: Search (RD) and Exhaustive (RD-EX)

RootDigger, introduced by [3], is a tool that takes an unrooted phylogenetic tree and its corresponding Multiple Sequence Alignment (MSA), and outputs a rooted tree. It uses a non-reversible Markov model to determine the most likely root location on the tree and infer confidence values for each potential root placement. It attempts to addresses the limitations of existing methods like molecular clock analysis (including midpoint rooting) and outgroup rooting. RootDigger circumvents the computationally demanding task of inferring a tree with a non-reversible model by using a reversible model for fast tree inference and then applying a non-reversible model solely for rooting the tree in a final step.

RootDigger was used in [26] for Phylogenetic analysis of bestrhodopsins. Here traditional outgroup method failed due to significant evolutionary distance. Alternative approaches such as RootDigger, which uses non-reversible Markov models, was applied to improve rooting accuracy.

RootDigger operates in two modes: Search and Exhaustive. The Search mode quickly identifies the most likely root using heuristics, while the Exhaustive mode thoroughly evaluates the likelihood of placing the root in every branch of the tree, reporting the Likelihood Weight Ratio for each branch. We considered both the search (RD) and exhaustive (RD-EX) modes for our study.

##### 2.5 Minimal Ancestor Deviation Rooting Method (MAD)

The Minimal Ancestor Deviation (MAD) rooting method is a phylogenetic technique designed to identify the root of an unrooted tree by minimizing deviations from the strict molecular clock hypothesis [33]. While strict ultrametricity rarely holds in practice, the midpoint criterion asserts that the middle of the path between two operational taxonomic units (OTUs) should coincide with their last common ancestor (LCA). The MAD algorithm evaluates this criterion by considering each branch of the tree as a potential root and then calculating the mean relative deviation from the molecular clock expectation for all OTU pairs. The branch that minimizes the deviation is considered the best candidate for the root. The deviation between the observed and expected distances from a putative ancestor node to any two OTUs is quantified by comparing the distance of each OTU to the ancestor with half the distance between the two OTUs. This method tries to ensure that the root placement is the one that best aligns with the expected molecular clock-based distances.

### 3 Supplementary Results and Discussion

#### 3.1 Supplementary Tables

Table S2: Average Triplet Score (TS)

| **Dataset** | **Model Cond$^n$** | **ASTRAL** | **STELAR** | | | | | | |
|---|---|---|---|---|---|---|---|---|---|
| | | | **MAD** | **MP** | **MV** | **OG** | **RAND** | **RD** | **RD-EX** |
| **11-taxon** | strong-5 | 761.65 | | 641.6 | 641.6 | 761.65 | 533.947 | | |
| | strong-15 | 2290.65 | | 1930.45 | 1930.45 | 2291.1 | 1851.1 | | |
| | strong-25 | 3798.6 | | 3198.6 | 3198.6 | 3798.6 | 3149.2 | | |
| | strong-50 | 7589.55 | | 6389.6 | 6389.6 | 7589.8 | 6352.3 | | |
| | strong-100 | 15164.4 | | 12764.4 | 12764.4 | 15165.05 | 12733.9 | | |
| **15-taxon** | 100-100 | 15184.2 | 17235.2 | 17695.4 | 17695.4 | 17695.4 | 16457.4 | 17675.7 | 16868.3 |
| | 100-1000 | 17272.8 | 20110.1 | 20290.5 | 20290.5 | 20290.5 | 18536.9 | 20340.2 | 19806.4 |
| | 100-true | 25885.8 | | 23959.9 | 22207.4 | 32822.3 | 19874.4 | | |
| | 1000-100 | 149866.4 | 171608.1 | 175925.6 | 175925.6 | 175925.6 | 163764.7 | 175895.2 | 167931.7 |
| | 1000-1000 | 170197.6 | 198719.1 | 199613.3 | 199613.3 | 199613.3 | 182300.7 | 201799.8 | 194817.4 |
| | 1000-true | 258798.3 | | 241057.2 | 218618.5 | 327645.8 | 203841.5 | | |
| **37-taxon** | 0.5X-200-500 | 928498.4 | 989474.5 | 989467.45 | 989148.2 | 989344.1 | 874149.65 | 843120.75 | 981842.95 |
| | 1X-200-1000 | 1051124.35 | 1158433.25 | 1158536.85 | 1158536.85 | 1158536.3 | 961302.25 | 940074.8 | 1138218.45 |
| | 1X-200-500 | 981359.05 | 1101439.55 | 1101722.65 | 1101557.0 | 1102434.0 | 952731.8 | 932451.85 | 1094015.15 |
| | 1X-200-true | 1305923.7 | 1410709.85 | 1410709.85 | 1410709.85 | 1410709.85 | 998847.45 | | |
| | 1X-400-1000 | 2218385.85 | 2314597.85 | 2314598.0 | 2314598.0 | 2314598.0 | 1926274.9 | 1925994.8 | 2275069.85 |
| | 1X-400-500 | 2001280.158 | 2202684.211 | 2202711.0 | 2202711.0 | 2202709.421 | 1905627.263 | 1905137.263 | 2184632.263 |
| | 1X-400-true | 2729554.05 | 2820623.75 | 2820623.75 | 2820623.75 | 2820623.75 | 1998009.45 | | |
| | 1X-800-1000 | 4285710.65 | 4624759.05 | 4624759.05 | 4624759.05 | 4624759.05 | 3855959.8 | 3855226.2 | 4546550.85 |
| | 1X-800-500 | 4004590.9 | 4415965.4 | 4415965.4 | 4415965.4 | 4415965.4 | 3810705.75 | 3809699.85 | 4376343.25 |
| | 1X-800-true | 5461561.3 | 5479549.1 | 5643918.0 | 5643918.0 | 5643918.0 | 3997968.1 | | |
| | 2X-200-500 | 1006160.45 | 1152943.65 | 1154764.9 | 1153979.25 | 1154896.6 | 988994.0 | 967021.4 | 1145624.8 |
| **48-taxon** | 0.5X-1000-500 | 9020299.15 | 9500062.25 | 10357359.35 | 9574800.2 | 10608691.5 | 7930725.65 | 7518799.15 | 10449514.2 |
| | 1X-100-500 | 1009324.85 | 720083.6 | 1068425.35 | 884224.4 | 1177023.25 | 852615.85 | 833157.05 | 1048816.6 |
| | 1X-1000-500 | 9855638.35 | 7606195.8 | 11379455.1 | 8782297.3 | 11723809.35 | 8499165.85 | 8171234.05 | 11123677.3 |
| | 1X-200-500 | 1991144.1 | 1500354.9 | 2213331.85 | 1745214.8 | 2352178.85 | 1712047.25 | 1649962.45 | 2203511.45 |
| | 1X-25-500 | 254762.85 | 196975.8 | 270455.75 | 219774.65 | 298140.6 | 215021.0 | 212741.2 | 253673.0 |
| | 1X-50-500 | 507800.25 | 365375.65 | 543951.15 | 453327.45 | 591600.8 | 427917.35 | 420172.9 | 517154.55 |
| | 1X-500-500 | 4920514.05 | 3797144.3 | 5693187.55 | 4361695.9 | 5862315.2 | 4311531.4 | 4099042.55 | 5528409.1 |
| | 2X-1000-500 | 10634954.65 | 6965546.05 | 11458453.75 | 7945845.2 | 11915250.75 | 8947757.9 | 8574869.6 | 10978527.2 |
| **200-taxon** | estimated | 717159608.4 | 755349429.111 | 765279555.556 | 749798311.556 | 759675755.889 | 785804351.333 | | |
| **Amniota** | Amino Acid | 25132 | | 30807 | 25073 | 38136 | 29755 | | |
| | Nucleotide | 31162 | | 34600 | 28448 | 42448 | 28447 | | |
| **Angiosperm** | Nuclear | 912705 | 1214602 | 1200657 | 1200657 | 1200657 | 867932 | | |

Table S3: True Triplet Scores

| **Dataset** | **Model Cond$^n$** | **ASTRAL** | **STELAR** | | | | | | |
|---|---|---|---|---|---|---|---|---|---|
| | | | MAD | MP | MV | OG | RAND | RD | RD-EX |
| **11-taxon** | strong-5 | 781.2 | | 660.35 | 660.35 | 780.4 | 558.4 | | |
| | strong-15 | 2404.05 | | 2044.9 | 2044.9 | 2407.3 | 1936.35 | | |
| | strong-25 | 4024.65 | | 3425.05 | 3425.05 | 4025.05 | 3329.9 | | |
| | strong-50 | 8057.45 | | 6859.25 | 6859.25 | 8050.7 | 6748.0 | | |
| | strong-100 | 16124.4 | | 13726.8 | 13726.8 | 16116.95 | 13601.95 | | |
| **15-taxon** | 100gt-100bp | 19589.4 | 30030.818 | 30461.9 | 30461.9 | 30461.9 | 20156.0 | 28224.7 | 21472.9 |
| | 100gt-1000bp | 21719.3 | 32766.9 | 32521.4 | 32521.4 | 32521.4 | 20864.3 | 26324.6 | 23831.8 |
| | 1000gt-100bp | 187521.0 | 321010.5 | 309185.1 | 309185.1 | 309185.1 | 198069.8 | 289730.0 | 217874.7 |
| | 1000gt-1000bp | 231557.5 | 327645.8 | 326824.3 | 326824.3 | 326824.3 | 199191.7 | 262838.1 | 231898.8 |
| **37-taxon** | 0.5X-200-500 | 1105674.15 | 1248086.15 | 1248052.1 | 1248244.05 | 1248109.45 | 920812.15 | 920358.95 | 1149700.0 |
| | 1X-200-1000 | 1201271.65 | 1410385.05 | 1410652.65 | 1410652.65 | 1410650.6 | 998723.7 | 832956.55 | 1281463.75 |
| | 1X-200-500 | 1145167.2 | 1410381.45 | 1410467.05 | 1410405.45 | 1409539.75 | 998074.45 | 869667.85 | 1291754.75 |
| | 1X-400-1000 | 2591563.25 | 2820598.0 | 2820594.95 | 2820594.95 | 2820594.95 | 1997631.55 | 1997081.55 | 2563032.95 |
| | 1X-400-500 | 2355862.105 | 2820118.789 | 2820230.579 | 2820230.579 | 2820224.316 | 1996615.684 | 1995856.0 | 2562692.579 |
| | 1X-800-1000 | 4774830.25 | 5479549.1 | 5479549.1 | 5479549.1 | 5479549.1 | 3881310.75 | 3147458.75 | 4978768.85 |
| | 1X-800-500 | 4600609.9 | 5479505.2 | 5479505.2 | 5479505.2 | 5479505.2 | 3881010.3 | 3371166.75 | 4978624.65 |
| | 2X-200-500 | 1174273.3 | 1497359.75 | 1497454.1 | 1498962.2 | 1496526.2 | 1034799.55 | 1034311.6 | 1357399.2 |
| 48-taxon | 0.5X-1000-500 | 9636236.65 | 10051174.3 | 11095538.9 | 10139002.25 | 11530857.55 | 8288721.45 | 7824656.45 | 11219496.35 |
| | 1X-100-500 | 1065149.15 | 732817.85 | 1127397.95 | 918274.75 | 1251100.15 | 897977.55 | 877085.75 | 1106691.05 |
| | 1X-1000-500 | 10462831.35 | 7840046.25 | 12066355.55 | 9109073.35 | 12591978.75 | 8972138.9 | 8617815.35 | 11802114.85 |
| | 1X-200-500 | 2100679.5 | 1538912.75 | 2338739.1 | 1812038.7 | 2506970.35 | 1803167.7 | 1736660.8 | 2326202.4 |
| | 1X-50-500 | 528423.65 | 372201.95 | 571365.9 | 469727.05 | 624282.25 | 449897.6 | 441395.05 | 543559.8 |
| | 1X-500-500 | 5208654.0 | 3909483.9 | 6035359.7 | 4527019.1 | 6280826.55 | 4547023.75 | 4318498.2 | 5869280.75 |
| | 2X-1000-500 | 11741419.55 | 7213265.45 | 12494254.9 | 8357037.45 | 13361902.55 | 9542505.2 | 9084815.15 | 11773397.15 |

Table S4: Average Quartet Scores

| Dataset | Model Cond$^n$ | ASTRAL | STELAR | | | | | | |
|---|---|---|---|---|---|---|---|---|---|
| | | | MAD | MP | MV | OG | RAND | RD | RD-EX |
| 11-taxon | strong-100 | 26705.4 | | 26705.4 | 26705.4 | 26702.7 | 26535.85 | | |
| | strong-15 | 4068.25 | | 4066.6 | 4066.6 | 4064.6 | 3970.7 | | |
| | strong-25 | 6695.4 | | 6695.15 | 6695.15 | 6695.15 | 6596.05 | | |
| | strong-5 | 1357.4 | | 1356.9 | 1356.9 | 1355.35 | 1318.9 | | |
| | strong-50 | 13383.1 | | 13383.05 | 13383.05 | 13380.1 | 13190.25 | | |
| 15-taxon | 100gt-100bp | 69933.8 | 69686.3 | 69276.5 | 69276.5 | 69276.5 | 68780.0 | 69545.4 | 68252.7 |
| | 100gt-1000bp | 82166.6 | 82105.2 | 82013.2 | 82013.2 | 82013.2 | 80610.5 | 81830.5 | 81233.9 |
| | 100gt-true | 84634.9 | | 84516.7 | 84570.0 | 84634.9 | 83301.7 | | |
| | 1000gt-100bp | 693656.1 | 691299.8 | 684572.4 | 684572.4 | 684572.4 | 682891.75 | 685192.0 | 687215.4 |
| | 1000gt-1000bp | 818022.2 | 817937.3 | 817532.111 | 817532.111 | 817532.111 | 811553.333 | 815358.222 | 811824.1 |
| | 1000gt-true | 844184.3 | | 844184.3 | 844184.3 | 844184.3 | 833455.6 | | |
| 37-taxon | 0.5X-200-500 | 10007425.55 | 10007255.25 | 10007260.7 | 10007291.35 | 10007287.65 | 9999226.7 | 9995322.45 | 10006075.75 |
| | 1X-200-1000 | 11586641.45 | 11586613.85 | 11586550.25 | 11586550.25 | 11586540.25 | 11581925.35 | 11578626.2 | 11586006.4 |
| | 1X-200-500 | 11270657.55 | 11270311.8 | 11270455.75 | 11270303.45 | 11270534.45 | 11262629.65 | 11257710.1 | 11269601.25 |
| | 1X-200-true | 11746203.5 | 11746194.95 | 11746198.85 | 11746198.85 | 11746198.85 | 11741292.0 | | |
| | 1X-400-1000 | 23159900.85 | 23159890.65 | 23159900.85 | 23159900.85 | 23159900.85 | 23148064.55 | 23144744.4 | 23159090.3 |
| | 1X-400-500 | 22530090.421 | 22529758.158 | 22530050.053 | 22530050.053 | 22529985.526 | 22518243.421 | 22502535.105 | 22528736.053 |
| | 1X-400-true | 23487558.65 | 23487558.65 | 23487558.65 | 23487558.65 | 23487558.65 | 23481670.65 | | |
| | 1X-800-1000 | 46342464.15 | 46342464.15 | 46342464.15 | 46342464.15 | 46342464.15 | 46329619.95 | 46307948.85 | 46342464.15 |
| | 1X-800-500 | 45096874.45 | 45096874.45 | 45096874.45 | 45096874.45 | 45096874.45 | 45083925.95 | 45052620.65 | 45096285.3 |
| | 1X-800-true | 45632640.45 | 45632640.45 | 45632640.45 | 45632640.45 | 45632640.45 | 45621015.4 | | |
| | 2X-200-500 | 11947266.5 | 11945264.3 | 11947184.75 | 11947089.25 | 11947252.75 | 11943092.85 | 11939246.25 | 11947046.35 |
| 48-taxon | 0.5X-1000-500 | 110583148.55 | 103668933.6 | 109362112.0 | 104025102.85 | 110383433.15 | 109578470.35 | 109734374.2 | 110314684.7 |
| | 1X-100-500 | 12383876.5 | 12025621.5 | 12138032.25 | 11964202.2 | 12318052.45 | 12207533.7 | 12179035.15 | 12204053.45 |
| | 1X-1000-500 | 123024287.2 | 119274515.2 | 121790825.05 | 114929114.55 | 122916649.8 | 121998800.7 | 122241813.9 | 122035658.75 |
| | 1X-200-500 | 24713063.7 | 23962846.1 | 24378714.6 | 23863357.3 | 24642898.2 | 24418292.65 | 24372977.5 | 24465457.3 |
| | 1X-25-500 | 3150240.45 | 3052890.9 | 3051834.85 | 3043066.7 | 3115783.05 | 3067542.55 | 3054728.5 | 3093191.25 |
| | 1X-50-500 | 6234610.35 | 6065409.95 | 6058419.0 | 6010449.1 | 6182000.55 | 6122005.95 | 6084794.85 | 6118501.55 |
| | 1X-500-500 | 61574240.65 | 59710450.05 | 60903919.2 | 58540593.6 | 61477542.2 | 60979019.25 | 61104007.6 | 61057573.4 |
| | 2X-1000-500 | 131631393.9 | 129156563.95 | 129905861.95 | 126831800.25 | 131546355.5 | 130155966.9 | 130740197.05 | 130252231.35 |
| 200-taxon | estimated | 47622712046.3 | 47536451073.2 | 47446554046.6 | 47519602308.4 | 47615850756.2 | 47390448761.7 | | |
| Amniota | Amino Acid | 83604 | | 82250 | 82250 | 82250 | 82077 | | |
| | Nucleotide | 97890 | | 97890 | 97890 | 97744 | 97881 | | |
| Angiosperm | Nuclear | 11553053 | 10963619 | 10936281 | 10936281 | 10936281 | 11210579 | | |

Table S5: True Quartet Scores

| Dataset | Model Cond$^n$ | ASTRAL | STELAR | | | | | | |
|---|---|---|---|---|---|---|---|---|---|
| | | | MAD | MP | MV | OG | RAND | RD | RD-EX |
| 11-taxon | strong-100 | 30708.3 | | 30708.3 | 30708.3 | 30663.25 | 30039.25 | | |
| | strong-15 | 4513.6 | | 4517.15 | 4517.15 | 4533.65 | 4273.95 | | |
| | strong-25 | 7622.3 | | 7625.5 | 7625.5 | 7625.5 | 7283.15 | | |
| | strong-5 | 1391.85 | | 1386.95 | 1386.95 | 1387.5 | 1371.4 | | |
| | strong-50 | 15322.95 | | 15327.75 | 15327.75 | 15282.75 | 14743.55 | | |
| 15-taxon | 100gt-100bp | 82158.6 | 83019.8 | 80525.5 | 80525.5 | 80525.5 | 78400.5 | 80608.9 | 78943.1 |
| | 100gt-1000bp | 84345.1 | 84545.7 | 84127.9 | 84127.9 | 84127.9 | 82597.5 | 83759.7 | 83369.4 |
| | 1000gt-100bp | 827970.1 | 8421420 | 816941.9 | 816941.9 | 816941.9 | 815221.4 | 808329.0 | 823085.8 |
| | 1000gt-1000bp | 843362.8 | 844184.3 | 843362.8 | 843362.8 | 843362.8 | 837052.9 | 839741.3 | 837294.5 |
| 37-taxon | 0.5X-200-500 | 10311630.05 | 10312053.55 | 10311354.45 | 10311638.25 | 10311920.2 | 10304742.0 | 10296194.95 | 10311070.2 |
| | 1X-200-1000 | 11743809.5 | 11743714.8 | 11744886.3 | 11744886.3 | 11744820.7 | 11739393.55 | 11739601.85 | 11743395.4 |
| | 1X-200-500 | 11739721.0 | 11740608.15 | 11740822.35 | 11740575.95 | 11740228.2 | 11734417.9 | 11729777.7 | 11739398.1 |
| | 1X-400-1000 | 23486728.75 | 23486827.8 | 23486728.75 | 23486728.75 | 23486728.75 | 23470981.1 | 23467004.0 | 23486217.7 |
| | 1X-400-500 | 23482457.0 | 23479373.16 | 23481661.05 | 23481661.05 | 23481461.05 | 23468128.37 | 23445782.8 | 23478993.4 |
| | 1X-800-1000 | 45632640.45 | 45632640.45 | 45632640.45 | 45632640.45 | 45632640.45 | 45619021.2 | 45608973.15 | 45632640.4 |
| | 1X-800-500 | 45631302.8 | 45631302.8 | 45631302.8 | 45631302.8 | 45631302.8 | 45610570.45 | 45596005.65 | 45629582.8 |
| | 2X-200-500 | 12565281.6 | 12560555.4 | 12567292.7 | 12568800.8 | 12565175.0 | 12559268.9 | 12547261.1 | 12568200.0 |
| 48-taxon | 0.5X-1000-500 | 119513731.6 | 110530209.8 | 117731361.0 | 110982521.8 | 119135686.95 | 118129439.45 | 118354949.2 | 119030503.3 |
| | 1X-100-500 | 13567423.65 | 13051253.55 | 13248847.0 | 12954507.65 | 13454157.6 | 13359236.1 | 13315353.65 | 13340893.3 |
| | 1X-1000-500 | 135985986.4 | 129816269.5 | 133918071.6 | 124794923.4 | 135761706.2 | 134327102.0 | 134657697.5 | 134271816.7 |
| | 1X-200-500 | 27170167.45 | 25974395.8 | 26665882.2 | 25855260.35 | 27003290.0 | 26767338.2 | 26695842.15 | 26745445.5 |
| | 1X-50-500 | 6776224.95 | 6549690.9 | 6565844.35 | 6482442.4 | 6708723.9 | 6668536.7 | 6611222.05 | 6653193.35 |
| | 1X-500-500 | 67891776.05 | 64911557.35 | 66810392.75 | 63519122.5 | 67684969.8 | 66998284.4 | 67178572.9 | 67077301.2 |
| | 2X-1000-500 | 148393885.35 | 143157302.05 | 144655982.55 | 139243603.85 | 148183540.35 | 145197013.0 | 146399056.5 | 145292248.1 |

Table S6: Percentage of correctly rooted gene trees

| Dataset | Model Cond$^n$ | STELAR | | | | | | |
|---|---|---|---|---|---|---|---|---|
| | | MAD | MP | MV | OG | RAND | RD | RD-EX |
| **11-taxon** | strong-5 | | 0.0 | 0.0 | 100.0 | 11.0 | 0.0 | |
| | strong-15 | | 0.0 | 0.0 | 100.0 | 8.9 | 0.0 | |
| | strong-25 | | 0.0 | 0.0 | 100.0 | 9.0 | 0.0 | |
| | strong-50 | | 0.0 | 0.0 | 100.0 | 9.9 | 0.0 | |
| | strong-100 | | 0.0 | 0.0 | 100.0 | 9.15 | 0.0 | |
| **15-taxon** | 100-100 | 19.5 | 100.0 | 100.0 | 100.0 | 7.0 | 5.9 | 3.8 |
| | 100-1000 | 85.4 | 100.0 | 100.0 | 100.0 | 7.7 | 0.0 | 6.0 |
| | 100-true | 0.0 | 0.0 | 0.0 | 100.0 | 6.5 | | |
| | 1000-100 | 19.6 | 100.0 | 100.0 | 100.0 | 5.7 | 0.5 | 3.9 |
| | 1000-1000 | 83.8 | 100.0 | 100.0 | 100.0 | 5.9 | 0.0 | 5.7 |
| | 1000-true | 0.0 | 0.0 | 0.0 | 100.0 | 6.1 | | |
| **37-taxon** | 0.5X-200-500 | 60.35 | 70.75 | 79.15 | 100.0 | 2.4 | 0.1 | 3.6 |
| | 1X-200-1000 | 78.75 | 81.35 | 86.9 | 100.0 | 2.4 | 0.1 | 5.0 |
| | 1X-200-500 | 66.85 | 77.6 | 85.35 | 100.0 | 2.5 | 0.0 | 5.45 |
| | 1X-400-1000 | 79.25 | 81.3 | 87.0 | 100.0 | 2.35 | 0.0 | 5.25 |
| | 1X-400-500 | 64.6 | 74.25 | 81.1 | 100.0 | 2.25 | 0.0 | 4.95 |
| | 1X-800-1000 | 78.5 | 80.9 | 86.55 | 100.0 | 2.45 | 0.0 | 5.1 |
| | 1X-800-500 | 67.45 | 78.2 | 85.25 | 100.0 | 2.3 | 0.0 | 5.3 |
| | 2X-200-500 | 71.6 | 80.8 | 87.1 | 100.0 | 2.8 | 0.1 | 6.35 |
| **48-taxon** | 0.5X-1000-500 | 7.0 | 0.6 | 6.55 | 100.0 | 1.7 | 0.0 | 0.2 |
| | 1X-100-500 | 6.2 | 1.0 | 7.25 | 100.0 | 1.8 | 0.0 | 0.0 |
| | 1X-1000-500 | 5.15 | 0.75 | 6.1 | 100.0 | 1.55 | 0.0 | 0.0 |
| | 1X-200-500 | 5.7 | 0.9 | 6.85 | 100.0 | 1.6 | 0.0 | 0.0 |
| | 1X-25-500 | 7.0 | 0.8 | 8.4 | 100.0 | 2.2 | 0.0 | 0.0 |
| | 1X-50-500 | 6.3 | 1.2 | 7.6 | 100.0 | 2.2 | 0.0 | 0.0 |
| | 1X-500-500 | 5.25 | 0.7 | 6.2 | 100.0 | 1.75 | 0.0 | 0.0 |
| | 2X-1000-500 | 3.6 | 0.8 | 5.0 | 100.0 | 1.7 | 0.0 | 0.0 |
| **200-taxon** | estimated | 6.6 | 9.3 | 11.2 | 100.0 | 0.0 | | |
| **500-taxon** | true | 25.3 | 0.0 | 0.0 | 100.0 | 0.0 | | |
| | estimated | 7.4 | 12.5 | 16.7 | 100.0 | 0.0 | | |

Table S7: Percentage of correctly rooted species trees

| Dataset | Model Cond$^n$ | STELAR | | | | | | |
|---|---|---|---|---|---|---|---|---|
| | | **MAD** | **MP** | **MV** | **OG** | **RAND** | **RD** | **RD-EX** |
| **11-taxon** | strong-5 | | 0.0 | 0.0 | 100.0 | 0.0 | 0.0 | |
| | strong-15 | | 0.0 | 0.0 | 100.0 | 0.0 | 0.0 | |
| | strong-25 | | 0.0 | 0.0 | 100.0 | 0.0 | 0.0 | |
| | strong-50 | | 0.0 | 0.0 | 100.0 | 0.0 | 0.0 | |
| | strong-100 | | 0.0 | 0.0 | 100.0 | 0.0 | 0.0 | |
| **15-taxon** | 100-100 | 50.0 | 100.0 | 100.0 | 100.0 | 0.0 | 60.0 | 0.0 |
| | 100-1000 | 100.0 | 100.0 | 100.0 | 100.0 | 0.0 | 0.0 | 0.0 |
| | 100-true | 0.0 | 0.0 | 0.0 | 100.0 | 0.0 | | |
| | 1000-100 | 80.0 | 100.0 | 100.0 | 100.0 | 0.0 | 90.0 | 0.0 |
| | 1000-1000 | 100.0 | 100.0 | 100.0 | 100.0 | 0.0 | 0.0 | 0.0 |
| | 1000-true | 0.0 | 0.0 | 0.0 | 100.0 | 0.0 | | |
| **37-taxon** | 0.5X-200-500 | 100.0 | 100.0 | 100.0 | 100.0 | 0.0 | 0.0 | 0.0 |
| | 1X-200-1000 | 100.0 | 100.0 | 100.0 | 100.0 | 0.0 | 0.0 | 0.0 |
| | 1X-200-500 | 100.0 | 100.0 | 100.0 | 100.0 | 0.0 | 0.0 | 0.0 |
| | 1X-400-1000 | 100.0 | 100.0 | 100.0 | 100.0 | 0.0 | 0.0 | 0.0 |
| | 1X-400-500 | 100.0 | 100.0 | 100.0 | 100.0 | 0.0 | 0.0 | 0.0 |
| | 1X-800-1000 | 100.0 | 100.0 | 100.0 | 100.0 | 0.0 | 0.0 | 0.0 |
| | 1X-800-500 | 100.0 | 100.0 | 100.0 | 100.0 | 0.0 | 0.0 | 0.0 |
| | 2X-200-500 | 100.0 | 100.0 | 100.0 | 100.0 | 0.0 | 0.0 | 0.0 |
| **48-taxon** | 0.5X-1000-500 | 0.0 | 0.0 | 0.0 | 100.0 | 0.0 | 0.0 | 0.0 |
| | 1X-100-500 | 0.0 | 0.0 | 0.0 | 100.0 | 0.0 | 0.0 | 0.0 |
| | 1X-1000-500 | 0.0 | 0.0 | 0.0 | 100.0 | 0.0 | 0.0 | 0.0 |
| | 1X-200-500 | 0.0 | 0.0 | 0.0 | 100.0 | 0.0 | 0.0 | 0.0 |
| | 1X-25-500 | 0.0 | 0.0 | 0.0 | 100.0 | 0.0 | 0.0 | 0.0 |
| | 1X-50-500 | 0.0 | 0.0 | 0.0 | 100.0 | 0.0 | 0.0 | 0.0 |
| | 1X-500-500 | 0.0 | 0.0 | 0.0 | 100.0 | 0.0 | 0.0 | 0.0 |
| | 2X-1000-500 | 0.0 | 0.0 | 0.0 | 100.0 | 0.0 | 0.0 | 0.0 |
| **200-taxon** | estimated | 30.0 | 40.0 | 40.0 | 100.0 | 0.0 | | |
| **500-taxon** | true | 40.0 | 0.0 | 0.0 | 100.0 | 0.0 | | |
| | estimated | 10.0 | 20.0 | 30.0 | 100.0 | 0.0 | | |

Table S8: Average Topological Distance between true and estimated roots of gene trees

| Dataset | Model Cond$^n$ | MAD | MP | MV | OG | RAND | RD | RD-EX |
|---|---|---|---|---|---|---|---|---|
| **11-taxon** | strong-5 | | 1.18 | 1.33 | 0.0 | 3.34 | | |
| | strong-15 | | 1.2 | 1.353 | 0.0 | 3.47 | | |
| | strong-25 | | 1.182 | 1.344 | 0.0 | 3.42 | | |
| | strong-50 | | 1.192 | 1.339 | 0.0 | 3.392 | | |
| | strong-100 | | 1.199 | 1.347 | 0.0 | 3.475 | | |
| **15-taxon** | 100gt-100bp | 3.536 | 0.0 | 0.0 | 0.0 | 5.253 | 3.685 | 5.343 |
| | 100gt-1000bp | 0.763 | 0.0 | 0.0 | 0.0 | 5.031 | 3.372 | 4.769 |
| | 100gt-true | | 2.948 | 3.159 | 0.0 | 5.048 | | |
| | 1000gt-100bp | 3.516 | 0.001 | 0.007 | 0.0 | 5.329 | 3.738 | 5.271 |
| | 1000gt-1000bp | 0.82 | 0.0 | 0.0 | 0.0 | 5.13 | 3.436 | 4.814 |
| | 1000gt-true | | 2.916 | 3.105 | 0.0 | 4.958 | | |
| **37-taxon** | 0.5X-200-500 | 1.423 | 0.634 | 0.547 | 0.0 | 7.606 | 8.085 | 4.378 |
| | 1X-200-1000 | 0.604 | 0.387 | 0.282 | 0.0 | 7.65 | 8.072 | 4.187 |
| | 1X-200-500 | 1.374 | 0.557 | 0.44 | 0.0 | 7.745 | 8.149 | 4.221 |
| | 1X-400-1000 | 0.602 | 0.391 | 0.282 | 0.0 | 7.616 | 8.094 | 4.172 |
| | 1X-400-500 | 1.341 | 0.551 | 0.444 | 0.0 | 7.649 | 8.168 | 4.224 |
| | 1X-800-1000 | 0.614 | 0.405 | 0.297 | 0.0 | 7.574 | 8.108 | 4.176 |
| | 1X-800-500 | 1.348 | 0.563 | 0.454 | 0.0 | 7.702 | 8.172 | 4.254 |
| | 2X-200-500 | 1.298 | 0.555 | 0.443 | 0.0 | 7.689 | 8.107 | 4.183 |
| **48-taxon** | 0.5X-1000-500 | 5.555 | 5.075 | 5.677 | 0.0 | 9.026 | 9.324 | 4.746 |
| | 1X-100-500 | 7.554 | 5.7 | 6.815 | 0.0 | 9.267 | 9.514 | 7.56 |
| | 1X-1000-500 | 7.523 | 5.65 | 6.861 | 0.0 | 9.247 | 9.538 | 7.563 |
| | 1X-200-500 | 7.497 | 5.637 | 6.809 | 0.0 | 9.187 | 9.472 | 7.516 |
| | 1X-25-500 | 7.034 | 5.386 | 6.402 | 0.0 | 9.158 | 9.304 | 7.452 |
| | 1X-50-500 | 7.301 | 5.449 | 6.643 | 0.0 | 9.116 | 9.391 | 7.501 |
| | 1X-500-500 | 7.495 | 5.614 | 6.832 | 0.0 | 9.121 | 9.545 | 7.548 |
| | 2X-1000-500 | 8.599 | 6.047 | 7.496 | 0.0 | 9.364 | 9.83 | 7.925 |
| **200-taxon** | estimated | 4.684 | 4.74 | 4.76 | 0.0 | 11.091 | | |
| **500-taxon** | true | 2.632 | 2.51 | 2.326 | 0.0 | 11.739 | | |
| | estimated | 5.612 | 5.917 | 5.875 | 0.0 | 14.15 | | |

Table S9: Average Topological Distance between true and estimated roots of species tree

| Dataset | Model Cond$^n$ | MAD | MP | MV | OG | RAND | RD | RD-EX |
|---|---|---|---|---|---|---|---|---|
| **11-taxon** | strong-5 | | 1.0 | 1.0 | 0.0 | 2.0 | | |
| | strong-15 | | 1.0 | 1.0 | 0.0 | 1.15 | | |
| | strong-25 | | 1.0 | 1.0 | 0.0 | 1.05 | | |
| | strong-50 | | 1.0 | 1.0 | 0.0 | 1.0 | | |
| | strong-100 | | 1.0 | 1.0 | 0.0 | 1.0 | | |
| **15-taxon** | 100gt-100bp | 1.1 | 0.0 | 0.0 | 0.0 | 2.8 | 0.4 | 3.6 |
| | 100gt-1000bp | 0.0 | 0.0 | 0.0 | 0.0 | 4.4 | 2.5 | 4.0 |
| | 100gt-true | | 4.9 | 5.6 | 0.0 | 6.0 | | |
| | 1000gt-100bp | 0.2 | 0.0 | 0.0 | 0.0 | 3.0 | 0.1 | 3.3 |
| | 1000gt-1000bp | 0.0 | 0.0 | 0.0 | 0.0 | 5.5 | 3.0 | 4.8 |
| | 1000gt-true | | 5.0 | 5.9 | 0.0 | 5.8 | | |
| **37-taxon** | 0.5X-200-500 | 0.0 | 0.0 | 0.0 | 0.0 | 4.1 | 4.45 | 2.85 |
| | 1X-200-1000 | 0.0 | 0.0 | 0.0 | 0.0 | 4.05 | 4.1 | 3.1 |
| | 1X-200-500 | 0.0 | 0.0 | 0.0 | 0.0 | 4.05 | 4.15 | 2.95 |
| | 1X-400-1000 | 0.0 | 0.0 | 0.0 | 0.0 | 4.0 | 4.05 | 3.0 |
| | 1X-400-500 | 0.0 | 0.0 | 0.0 | 0.0 | 4.053 | 4.053 | 3.0 |
| | 1X-800-1000 | 0.0 | 0.0 | 0.0 | 0.0 | 4.0 | 4.0 | 3.0 |
| | 1X-800-500 | 0.0 | 0.0 | 0.0 | 0.0 | 4.0 | 4.0 | 3.0 |
| | 2X-200-500 | 0.0 | 0.0 | 0.0 | 0.0 | 3.95 | 4.1 | 2.9 |
| **48-taxon** | 0.5X-1000-500 | 2.0 | 2.0 | 2.0 | 0.0 | 4.7 | 8.6 | 2.0 |
| | 1X-100-500 | 5.65 | 2.8 | 4.0 | 0.0 | 6.0 | 8.0 | 3.4 |
| | 1X-1000-500 | 4.05 | 2.0 | 3.2 | 0.0 | 5.0 | 8.6 | 3.0 |
| | 1X-200-500 | 4.7 | 2.35 | 3.8 | 0.0 | 5.5 | 8.4 | 3.15 |
| | 1X-25-500 | 6.2 | 2.7 | 4.6 | 0.0 | 6.75 | 7.0 | 4.6 |
| | 1X-50-500 | 6.05 | 2.55 | 4.0 | 0.0 | 5.95 | 7.25 | 4.25 |
| | 1X-500-500 | 4.15 | 2.0 | 3.35 | 0.0 | 4.8 | 8.35 | 3.0 |
| | 2X-1000-500 | 5.0 | 2.05 | 3.9 | 0.0 | 4.7 | 8.85 | 3.35 |
| **200-taxon** | estimated | 1.0 | 1.0 | 1.1 | 0.0 | 3.5 | | |
| **500-taxon** | true | 3.1 | 2.2 | 2.5 | 0.0 | 6.4 | | |
| | estimated | 3.444 | 4.9 | 3.4 | 0.0 | 7.0 | | |

##### 3.2 Results on 11 Taxa Dataset

With varying genecount, the RF scores (figure S1) depict a general trend similar to that of the 37-taxa and 15-taxa datasets: increasing genecounts results in more accurate species trees for all methods, leading to lower RF scores. All variants of STELAR (except STELAR-RAND) yielded almost identical performances in terms of RF scores when compared to ASTRAL, with STELAR-MP and STELAR-MV yielding the same species trees in all cases.

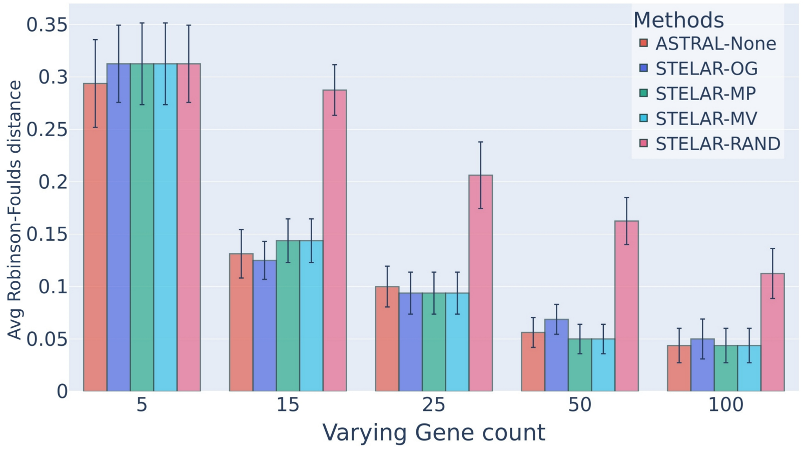


Fig. S1: Comparison of the Robinson-Fould's distance for ATRRAL and the STELAR variants on the 11 taxa dataset. We varied the gene counts from 5 to 100, considering only the high ILS condition.

Compared to the previous datasets, the differences in RF scores were more subtle in this case. These could not be captured completely by the triplet scores, for example - despite having a higher triplet score than ASTRAL in the 50 and 100 gene conditions (Table S2), STELAR-OG had a slightly worse RF score (figure S1). Upon inspecting the True Triplet Scores (TTS) from Table S3, we find that a worse RF score actually corresponded to a worse TTS for STELAR-OG. Other model conditions also mirrored this trend, where the TTS (and not the Triplet Scores) were a good indicator of the RF scores.

Referring to the Quartet Scores in Table S4, we see that ASTRAL has marginally higher quartet scores in each case despite not always being the best in terms of RF scores, a trend that carries over from prior datasets. Similar to the TTS, we find that True Quartet Scores (TQS) from Table S5 better capture the subtle changes in RF scores.

Referring to Table S8, we see that MV displayed a slightly higher inaccuracy in rooting the gene trees, when compared to MP. Despite this, STELAR still maintains its robustness in estimating the rooting of the final species tree in both cases (Table S9), with the estimated root differing from the true root by one node on average.

##### 3.3 Results on 48 Taxa Dataset

In line with our observations thus far, all methods followed the trend of better RF scores with "easier" model conditions (Figure S2). Unlike thus far, STELAR yielded better results with outgroup rooting in this particular dataset. Despite yielding competitive performances in "harder" conditions, STELAR-MP, STELAR-MV, STELAR-MAD failed to keep pace with STELAR-OG in higher gene counts (Figure S2(a)) and lower ILS (Figure S2(b)). Furthermore, STELAR proved to be more robust to random rooting - yielding the best performance by STELAR-RAND so far. This reinforces the prior hypothesis of RAND rooting performing better as taxa count increases. We also see that both STELAR-RD and STELAR-RD-EX yielded rather comparable results in most model conditions.

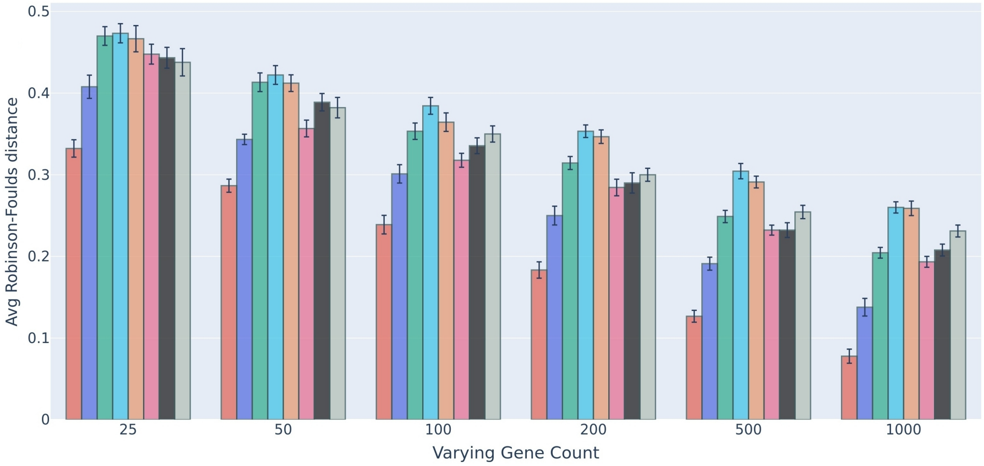


(a)

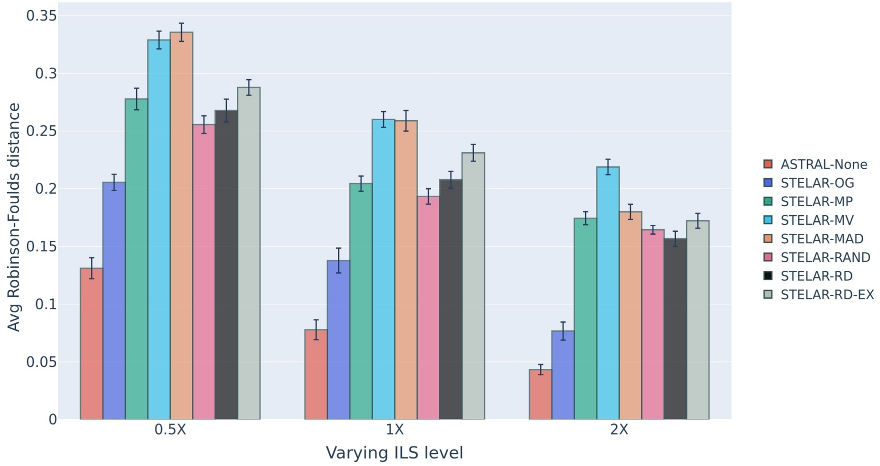


(b)

Fig. S2: Comparison of Average Robinson-Foulds distance for ASTRAL and STELAR with different rooting methods for the 48-taxa avian dataset. (a) We fixed the ILS at moderate (1X) and the base pairs at 500bp, and varied gene counts between from 25 to 1000. (b) We set the gene trees at 1000, the sequence length at 500bp, and varied the ILS between high (0.5X), moderate (1X) and low (2X)

As with previous observations, the Triplet Score followed the trend of increasing with lower RF scores and vice versa (Table S2). However, exceptions were observed in this dataset for STELAR-RAND and STELAR-RD. These exceptions persisted even when looking at the True Triplet Scores (Table S3), similar to observations in the 37 taxa dataset. On the other hand, Quartet scores (Table S4 were a better indicator of RF-scores in this dataset - modeling the trend that “higher quartet scores mean lower RF scores” better than previous datasets. The few discrepancies in this trend are all solved if we look at the True Quartet Scores (TQS) from Table: S5, where a higher TQS almost always means a lower RF score, which was not always the case in the 37-taxa dataset.

In order to explain the cause for the decline in performance for STELAR-MP, STELAR-MV and STELAR-MAD, we inspect the quality of gene tree rooting depicted in Table S8. Compared to the previous datasets, we find gross inaccuracies in gene tree rooting for MV and MAD, as well as for MP in a slightly less severe sense. Since the rooting methods MP, MV, and MAD are closely dependent on the branch lengths and branch supports of the input gene trees, we suspect irregularities in these values for this 48-taxa dataset, leading to worse gene tree rootings. This irregularity in gene tree rooting led to worse performance for the corresponding STELAR variants. Similar to 15 and 37 taxa datasets, gene tree rootings by RD and RAND had very high topological distances from the true root, leading to lower TS and TTS. However, unlike before, their RF scores were indeed competitive this time, as discussed above. Furthermore, unlike before, STELAR had a difficult time predicting the root this time, as indicated by none of the variants correctly predicting the root (Table S7) and the high topological distances between species tree root and true root (Table S8).
